# *In vivo* engineered quadrivalent CAR-macrophages break tumor barrier, remodel TME and overcome tumor heterogeneity through antigen spreading

**DOI:** 10.1101/2024.11.06.621821

**Authors:** Shaolong Zhang, Hengxing Lu, Hailing Zhang, Xizhong Ding, Qixin Wu, Anhua Lei, Na Kong, Jin Zhang

## Abstract

CAR-macrophages have shown promising prospect in treating solid tumors. *Ex vivo* engineered CAR-macrophages face the challenge of low transfection efficiency, long preparation cycle, and limited cell number. Here, we designed two novel CAR molecules, GPC3-CAR-Super IL-2 and FAP-CAR-△TGFβRII from the perspective of targeting tumor cells and improving tumor microenvironment, and generated quadrivalent CAR-macrophages *in vivo* for treating solid tumors by the LNP-mRNA system. We found that *in vivo* engineered quadrivalent CAR-macrophages can strongly activate tumor immunity and achieve complete tumor regression without significant side effects. Mechanically, *in vivo* engineered quadrivalent CAR-macrophages broke down physical barriers around the tumor constructed by CAFs and significantly promoted infiltration and expansion of CD8^+^ T cells. Moreover, the transiently formed CAR-macrophages *in vivo* are sufficient to form long-lasting T cell memory which can effectively prevent tumor recurrence. Most importantly, *in vivo* engineered CAR-macrophages also stimulated T cell memory against antigen-negative tumor cells through antigen spreading, which might effectively prevent the immune escape of heterogeneous tumor cells. Overall, we developed a platform of *in vivo* CAR-macrophages with dual roles as a tumor-killing effector cell and a recurrence-preventing vaccine.

## Introduction

Chimeric antigen receptor (CAR) T cell therapies have achieved remarkable success in the treatment of hematologic malignancies. However, CAR-T cells have so far shown limited efficacy against solid tumors, mainly due to the low infiltration and T cell exhaustion caused by the immunosuppressive tumor microenvironment (TME)[1]. In this context, CAR-macrophages have emerged to treat solid tumors due to their high infiltration, phagocytosis activity, and capacity to regulate the immune microenvironment[2, 3]. However, *ex vivo* engineered CAR-macrophages are faced with problems such as low transfection efficiency, limited cell number, and long preparation cycles. These challenges greatly limit the clinical application of CAR-macrophages in disease treatment. The successful development of mRNA vaccines against SARS-CoV-2 has validated the safety and efficacy of lipid nanoparticle (LNP) delivery. At present, the application of LNP-mRNA is not limited to vaccines in infectious diseases but has been extended to the field of gene editing and immune therapies[4]. LNPs were preferably enriched in the liver. Researchers have successfully achieved gene editing in liver macrophages by delivering Cre mRNA through LNP[5]. In addition, a CD5 monoclonal antibody-modified LNP delivery system was used to successfully generate FAP-targeting CAR-T cells *in vivo* to treat cardiac injury[6]. Based on these studies, we engineered the CAR-macrophages *in vivo* by the LNP-mRNA system for treating solid tumors.

The TME is a highly complex ecosystem that includes a variety of immune cells, cancer-associated fibroblasts (CAFs), endothelial cells, and extracellular matrix (ECM)[7]. CAFs are one of the most important components of the tumor microenvironment and play an essential role in the development of tumors. Activated CAFs can directly promote tumor growth and angiogenesis by secreting VEGF, and inhibit the immune response by secreting immunosuppressive cytokines[8]. Moreover, activated CAFs can create a physical barrier by remodeling ECM to inhibit the infiltration of drugs or immune cells[9]. In addition to physical barriers, CAFs can limit the infiltration of CD8^+^ T cells by establishing biochemical barriers. CXCL12 secreted by CAFs can interact with the CXCR4 receptor on the surface of CD8^+^ T cells to inhibit the infiltration of CD8^+^ T cells into the interior of the tumor[10]. TGFβ is the master regulator of CAF activation, which could maintain the continuous activation of CAFs through positive feedback[11]. In addition, TGFβ is also a well-known immunosuppressive cytokine and can inhibit macrophage phagocytosis, antigen presentation, and expression of inflammatory factors[12]. Therefore, minimizing TGF β signaling while simultaneously targeting CAFs by CAR-macrophages would be more helpful for treating solid tumors.

As a component of innate immunity, macrophages play an important role in shaping the functions of T cells. Numerous studies have shown spatiotemporal co-dependency between TAMs and exhausted CD8^+^ T cells in solid tumors[13–15]. TAMs could affect CD8+ T cell immunity by secreting cytokines or metabolic and epigenetic reprogramming. More recently it has been reported that TAMs restrict CD8^+^ T cell function through collagen deposition directed by TGFβ of the breast cancer microenvironment[16]. This further highlights the pivotal role of TAMs in fibrosis and CD8^+^ T cell exhaustion in solid tumors. Therefore, *in situ-*engineered TAMs may be an effective strategy to reactivate CD8^+^ T cells. Glypican-3 (GPC3) is specifically highly expressed in various solid tumors and is an ideal target for treating solid tumors[17]. We previously reported the anti-tumor effects of GPC3-CAR macrophages and their ability to activate CD8^+^ T cells through antigen presentation[18]. However, the complete activation of CD8^+^ T also requires cytokine stimulation. Among them, IL-2 is the most important cytokine that can promote the proliferation and activation of CD8^+^ T cells. However, natural IL-2 preferentially activates Treg cells that express high CD25 (IL-2Rα) rather than CD8^+^ T cells [19]. Recently, modified Super IL-2 not only showed a higher affinity for the CD122 (IL-2Rβ) receptor on the surface of CD8^+^ T cells (not Tregs) but also had better stability to specifically activate CD8^+^ T cells[20]. Therefore, *in vivo*-engineered CAR-macrophages that can secrete Super IL-2 *in situ* may be more conducive to fully activating CD8^+^ T cells.

Here we designed two novel CAR molecules: GPC3-CAR-Super IL-2 and FAP-CAR-△TGFβRII. They were delivered simultaneously by the LNP-mRNA system (LNP-GF CAR mRNA) to achieve *in vivo* editing of CAR-macrophages. *In vivo* engineered CAR-macrophages by LNP-GF CAR mRNA caused complete regression of orthotopic hepatocellular carcinoma (HCC) tumors. Moreover, they effectively broke the physical barrier constructed by CAFs and significantly promoted the infiltration, expansion, and activation of CD8^+^ T cells. Interestingly, we found that the transient formation of CAR-macrophages *in vivo* is sufficient to form T cell immune memory. Furthermore, we confirmed that *in vivo* engineered CAR-macrophages could promote immune memory formation against antigen-negative tumor cells through antigen spreading, which led the way to solving tumor cell escape due to the heterogeneity.

## Results

### Transplanted macrophages did not effectively infiltrate into solid tumors

The highly efficient infiltration of CAR-macrophages into solid tumors is the basis for its anti-tumor effect. However, systematic analysis of CAR-macrophage distribution within tumors *in vivo* has yet to be reported. Therefore, we investigated the macrophage distribution in a tumor model with intravenous injection of bone marrow-derived macrophages (BMDMs) of mice. Firstly we established a subcutaneous tumor model with Hepa1-6 cells. Two weeks later we injected DiD-labeled BMDMs intravenously into the mice. We found that the transplanted macrophages were mainly concentrated in the liver rather than the tumor (Figure 1A). This was confirmed by immunofluorescence analysis of the liver and tumor (Extended Data Fig. 1A). We speculate that due to the superior blood supply of the liver, the intravenously transplanted macrophages may have better infiltration in an orthotopic HCC tumor model. Thus, we established such a model by intrahepatic injection of Hepa1-6 cells. After the orthotopic HCC tumor model was established by live animal imaging, the DiD-labeled BMDMs were injected into these mice intravenously. Surprisingly, we found that the DiD signals were enriched in the normal liver tissue area outside the tumor (Figure 1B). Immunofluorescence analysis further verified this phenomenon, and the DiD-labeled BMDMs were strictly confined to the periphery of the tumor and could not infiltrate into the tumor interior (Figure 1C, Extended Data Fig. 1B). These results suggest that exogenous macrophages cannot infiltrate into solid tumors effectively, which might reduce their therapeutic efficacy and limit their clinical applications. We also examined the distribution of endogenous macrophages in the same liver cancer model, and immunofluorescence showed that they were widely distributed inside and outside the tumor (Figure 1D). Combining the results above, we summarized the problems faced by transplanting *ex vivo* engineered CAR-macrophages and decided to focus on the endogenous macrophage to engineer the CAR-macrophages *in situ* by the LNP-mRNA system to treat solid tumors (Figure 1E).

**Figure 1:**
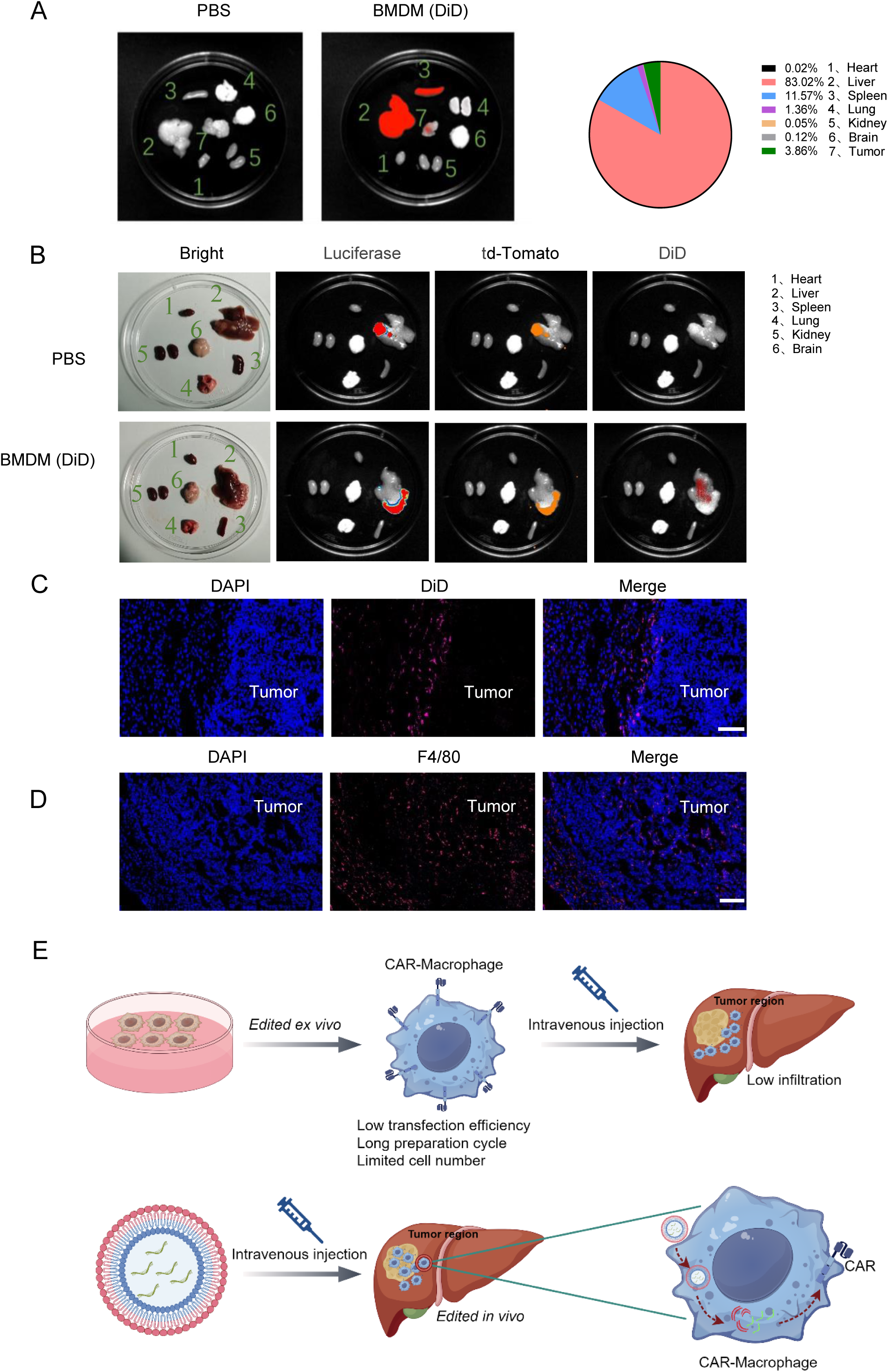
Transplanted macrophages did not effectively infiltrate into solid tumors. A: The subcutaneous tumor of Hepa1-6 cells was established in C57 mice. 2×10^6^ DiD-labeled BMDMs were injected intravenically and the tissues were collected for imaging after 3 days. The right panel is the DiD signal statistics. B: Orthotopic HCC tumor of Hepa1-6 (Luciferase^+^td-Tomato^+^) was established in C57 mice. 2×10^6^ DiD-labeled BMDMs were injected intravenically and the tissues were collected for imaging after 3 days. Luciferase and td-Tomato signals indicate the tumor area, and the DiD signal indicates the location of BMDMs. C: Immunofluorescence assays showing the distribution of DiD-labeled BMDMs in the liver of orthotopic HCC tumors. DiD signal (pink) indicated the location of BMDMs. Scale bar: 100 μm. D: Immunofluorescence staining of F4/80 (Red) in the liver tissue sections of orthotopic HCC tumor mice. Scale bar: 100 μm. E: A cartoon diagram illustrating the obstacle of *ex vivo* engineered CAR-macrophages in the treatment of solid tumors, and the proposed *in vivo* engineered CAR-macrophages strategy (By Figdraw).

### Design CAR molecules in the perspective of improving tumor microenvironment

In designing the mRNA molecules for *in vivo* engineered macrophages, we take full consideration of the tumor microenvironment. Previous spatial transcriptome data of hepatocellular carcinoma patients showed that there was a dense fibrous cell band composed of CAFs between the tumor area and the healthy tissue, and immune cells were confined to the periphery of the tumor by the CAFs[21, 22]. Fibroblast activation protein-α (FAP) is a typical marker of CAFs and is essential for ECM remodeling[23]. In addition, The Cancer Genome Atlas (TCGA) data showed that the expression level of FAP was negatively correlated with the survival rate of HCC patients (Figure 2A). Thus we speculated that the dense fibrocyte layer around the tumor may be the reason that transplanted macrophages were not able to infiltrate. Immunofluorescence staining of FAP with liver tissues confirmed the distribution of CAFs around the tumor (Figure 2B). The DiD-labeled macrophages were in close contact with FAP-positive cells but were strictly confined to the tumor periphery. Besides, CD8^+^ T cells were also strictly confined to the stroma region at the junction of the tumor and the healthy tissue (Figure 2C). These findings implied that breaking down the physical barrier constructed by CAFs is necessary to unleash the anti-tumor capacity of immune cells.

**Figure 2:**
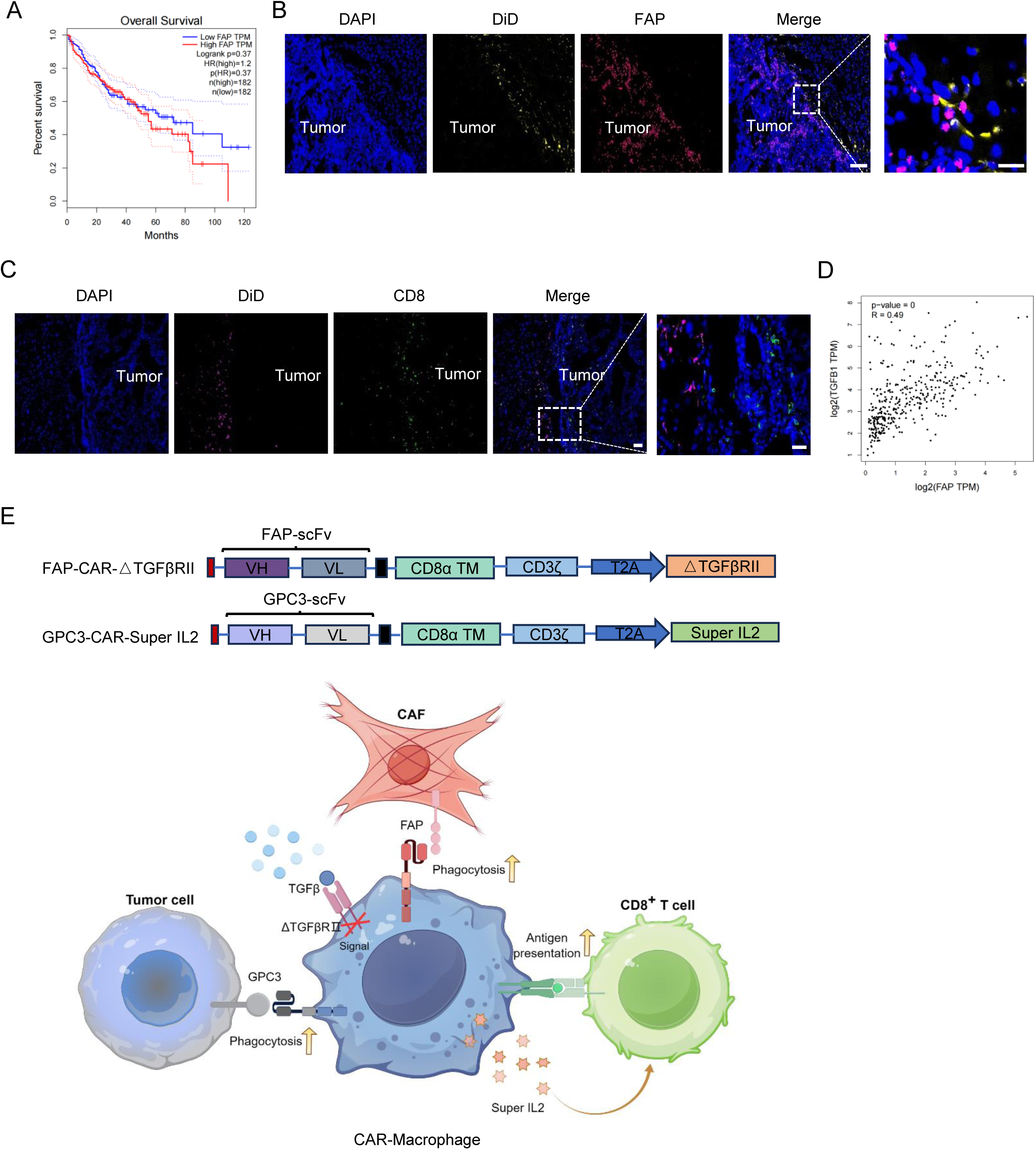
Design principle of CAR molecular structure. A: Kaplan–Meier curves of the survival analysis of HCC patients with high expression of FAP based on the TCGA database. B: Immunofluorescence assays showing the distribution of DiD-labeled BMDMs and CAFs in the liver of orthotopic HCC tumor mice. DiD (yellow) marked BMDMs and FAP (red) marked CAFs. Scale bar: 100 μm, magnified partial view Scale bar: 20 μm. C: Immunofluorescence staining of CD8 (Green) and DiD (pink) in the liver of orthotopic HCC tumor mice. Scale bar: 300 μm. D: Correlation analysis between FAP and TGFβ protein expression of HCC patients based on TCGA database. E: A cartoon diagram illustrating the design principle of the CAR molecules (By Figdraw).

Therefore, we designed a FAP-CAR for the *in vivo* engineered macrophages to target CAFs. Moreover, TGFβ is the core cytokine to activate CAFs, and it is also a well-known immunosuppressive cytokine, which can promote M2 polarization of macrophages[11, 12]. Clinical data from the TCGA database also showed that there was a significant positive correlation between the expression levels of FAP and TGFβ in hepatocellular carcinoma patients (Figure 2D). Studies have shown that the expression of an intracellular domain negative TGFβRII receptor (△TGFβRII) in cells can significantly inhibit the TGFβ signal[24, 25]. Therefore, we designed a FAP-CAR containing a dominant-negative TGFBRII (FAP-CAR-△TGFβRII), whose expression can block the TGFβ signaling (Figure 2E). We speculate that with the removal of the physical barrier, CD8^+^ T cells might be attracted from the stroma region, and get reactivated in the tumor. To confer the *in vivo* engineered CAR-macrophages with antigen specificity, we designed a GPC3 CAR for targeting liver cancer cells. As mentioned before, Super IL-2 was added to obtain a GPC3-CAR-Super IL-2 that could preferentially activate CD8^+^ T cells but not Treg cells (Figure 2E).

### LNP-GF CAR mRNA engineered CAR-macrophages showed dual-targeted phagocytosis, tolerance to TGFβ stimulation, and promotion of CD8^+^ T cell proliferation

Combining these designs, we aimed to edit CAR-macrophages *in vivo* to eliminate both tumor cells and CAFs, as well as unleash the anti-tumor capacity of CD8^+^ T cells. Ultimately, the final product of our LNP-mRNA system is to simultaneously deliver two mRNA molecules: FAP-CAR-△TGFβRII mRNA and GPC3-CAR-Super IL-2 mRNA. The mRNAs were transcribed *in vitro*, and LNPs were prepared by mixing the two mRNA components at a ratio of 1:1, which was referred to as LNP-GF CAR mRNA (Figure 3A). GFP mRNA was used as Control mRNA. To characterize the LNPs, we found the size of the LNPs was ∼150nm, the Polydispersity Index (PDI) was ∼0.1, and the electrical potential was ∼-3mV (Figure 3B). Next, we transfected BMDMs with the LNP-GF CAR mRNA, and found high expression of mRNAs of FAP-CAR, GPC3-CAR, △TGFβRII and Super IL-2, even though the levels declined overtime (Extended Data Fig. 2A). We also detected the expression of GPC3 and FAP CAR molecules by flow cytometry (Extended Data Fig. 2B). Together, these data indicate that we achieved dual-targeted CAR-macrophages editing with the LNP-mRNA system.

**Figure 3:**
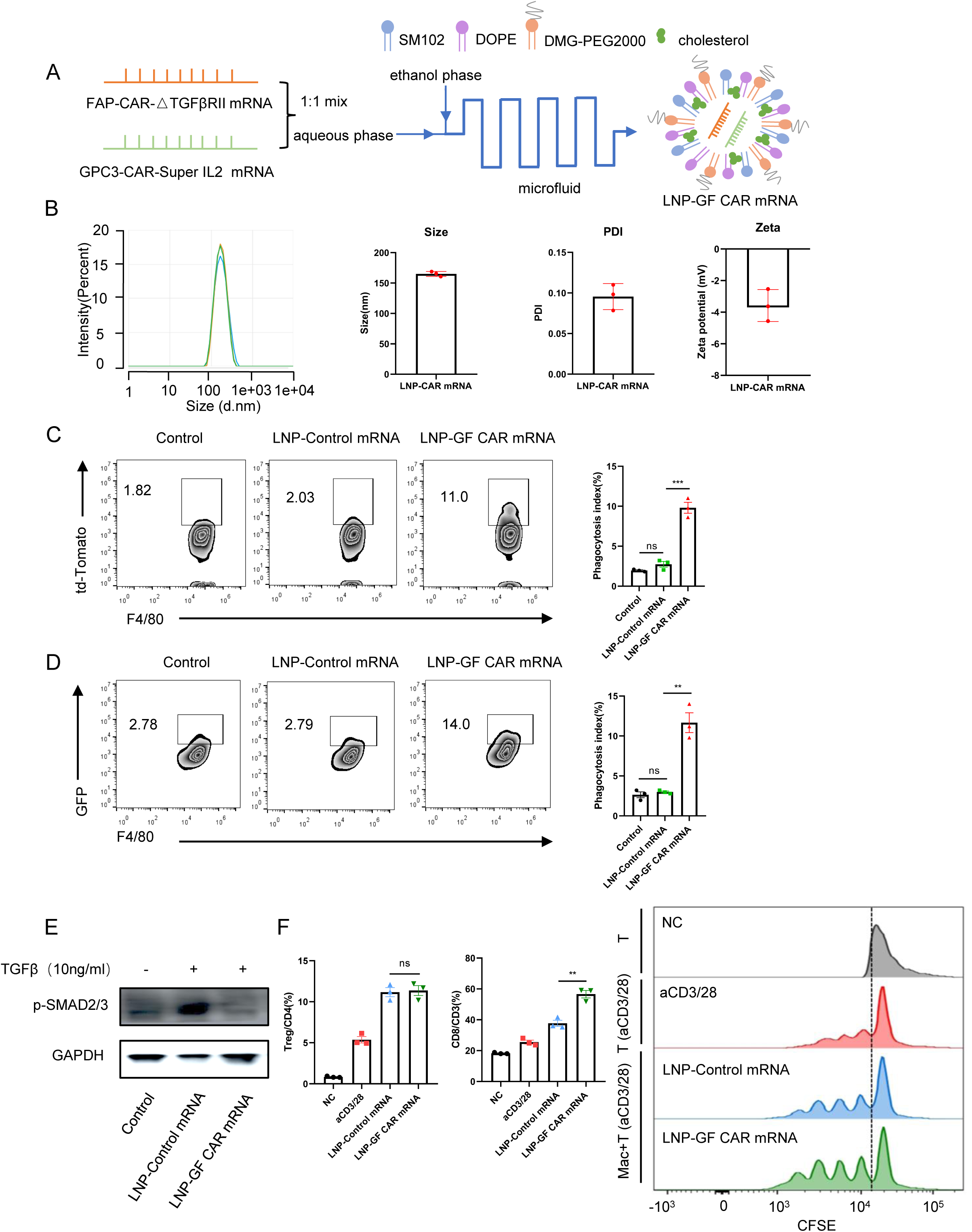
*Ex vivo* engineered CAR macrophages by LNP-GF CAR mRNA achieved the designed functions. A: A cartoon diagram illustrating the preparation process of LNP-GF CAR mRNA. B: Characterization of LNP-GF CAR mRNA (Hydrodynamic diameter, PDI, zeta potential, n = 3). C, D: BMDMs were transfected with LNP-Control mRNA or LNP-GF CAR mRNA (1ug/ml) for 24 hours. Engineered macrophages were co-incubated with GPC3^+^td-Tomato^+^Hepa1-6 cells and FAP^+^GFP^+^3T3 cells at an E/T ratio of 1:2 respectively. The phagocytosis of macrophages was detected after 48 hours of co-incubation by flow cytometry. Luciferase mRNA was used as the Control mRNA in this experiment. E: BMDMs were transfected with LNP-Control mRNA or LNP-GF CAR mRNA (1ug/ml) and TGFβ (10ng/ml) was added 24 hours later. The activation of p-SMAD2/3 was detected by western blotting after 30 minutes. F: BMDMs were transfected with LNP-Control mRNA or LNP-GF CAR mRNA (1ug/ml) for 24 hours and subsequently co-incubated with spleen T cells at a ratio of 1:5. After 3 days of co-incubation, the ratio of Treg and CD8^+^ T cells and the proliferation of CD8^+^ T cells indicated by CFSE staining were determined by flow cytometry. NC: Unstimulated T cells, aCD3/28: aCD3/28 stimulated T cells, LNP-Control mRNA: LNP-Control mRNA transfected macrophages, LNP-GF CAR mRNA: LNP-GF CAR mRNA transfected macrophages. Data represent the mean ± s.e.m. from a representative experiment (*n* = 3 biologically independent samples) of two independent experiments. Significance was calculated with one-way ANOVA analysis with multiple comparisons, ns: not significant; \*\**P* < 0.01; ***p < 0.001, \*\*\*\**P* < 0.0001.

Next, we explored whether CAR-macrophages engineered by LNP-GF CAR mRNA could achieve the designed functions such as dual-targeted phagocytosis. The LNP-GF CAR mRNA or LNP-Control mRNA was transfected into BMDMs. After 24 hours, macrophages engineered by the LNPs were co-incubated with Hepa1-6 cells expressing human GPC3 or 3T3 cells expressing mouse FAP, respectively. The results showed that compared with the LNP-Control mRNA, CAR-macrophages engineered by the LNP-GF CAR mRNA significantly increased the phagocytic activity against the GPC3^+^ Hepa1-6 cells and FAP^+^ 3T3 cells (Figure 3C, D). Next, we examined whether the CAR-macrophages engineered by LNP-GF CAR mRNA had reduced TGFβ signaling. The results showed that compared with the LNP-Control mRNA, CAR-macrophages engineered by the LNP-GF CAR mRNA almost completely abolished the activation of TGFβ downstream signaling indicated by phosphorylation of SMAD2/3 (Figure 3E). Moreover, we explored whether CAR-macrophages engineered by LNP-GF CAR mRNA could specifically promote proliferation of CD8^+^ T cells, by co-incubating the engineered CAR-macrophages with mouse spleen T cells. Results showed that CAR-macrophages engineered by the LNP-GF CAR mRNA did not affect the proportion of Treg cells, but significantly increased the proportion of CD8^+^ T cells and promoted the proliferation of CD8^+^ T cells indicated by the CFSE analysis (Figure 3F). Together, these results demonstrated that the CAR-macrophages engineered by LNP-GF CAR mRNA not only had the dual-targeted phagocytic capacity through the GPC3 and FAP CAR molecules, but also could tolerate TGFβ stimulation, and specifically promoted the proliferation of CD8^+^ T cells, which is defined as quadrivalent CAR macrophages.

### LNP-GF CAR mRNA engineered CAR-macrophages significantly inhibited the growth of tumor cells

We further explored the killing effect of LNP-GF CAR mRNA engineered CAR-macrophages against tumor cells. We transfected BMDMs with LNP-GF CAR mRNA or LNP-Control mRNA for 24 hours. The macrophages engineered by the LNPs were cocultured with Hepa1-6 cells (luciferase^+^GPC3^+^) with different effect/target ratios (E/T) of 2/1, 5/1, and 10/1. After 48 h of coculturing, the luciferase activity of the tumor cells was measured. We found that both a low and a high dose of LNP-GF CAR mRNA engineered CAR-macrophages could strongly inhibit tumor growth (Figure 4A). At an E/T ratio of 10:1, LNP-GF CAR mRNA engineered CAR-macrophages almost completely inhibited the growth of tumor cells. Even at an E/T ratio of 2:1, the killing rate was still about 60%. Next, we further explored the dynamic killing process by live imaging using an E/T ratio of 10:1. Consistent with previous results, CAR-macrophages engineered with all three doses of LNP-GF CAR mRNA could completely inhibit the growth of tumor cells (Figure 4B). Imaging data from Incucyte at the endpoint of 72 hours of co-incubation showed nearly no fluorescent signal in the LNP-GF CAR mRNA treated group (Extended Data Fig. 3A). With fluorescence microscopy performed 48 hours after coculture, we found that macrophages in the untransfected control or transfected with LNP-Control mRNA had almost no phagocytosis of tumor cells, whereas, in the LNP-GF CAR mRNA transfected group, the fluorescence signals of tumor cells were significantly reduced and fragmented (Figure 4C), suggesting they increased phagocytosis of CAR-macrophages against tumor cells. To dissect the individual role of the FAP-CAR-△TGFβRII mRNA versus the GPC3-CAR-Super IL-2 mRNA, we used each of them separately for the killing experiment on tumor cells again. The results showed that compared with the LNP-Control mRNA, CAR-macrophages engineered by LNP-FAP-CAR-△TGFβRII mRNA had no killing capacity against the GPC3^+^ Hepa1-6 cells. In contrast, the LNP-GPC3-CAR-Super IL-2 mRNA showed a similar killing effect compared with the LNP-GF CAR mRNA on GPC3^+^ Hepa1-6 cells (Figure 4D), indicating that the killing effect of LNP-GF CAR mRNA was attributable to the GPC3-CAR. Next, we examined whether the killing of target cells by CAR-macrophages was antigen specific. LNP-GF CAR mRNA engineered CAR-macrophages were incubated with td-Tomato^+^ WT Hepa1-6 cells (GPC3 negative) and td-Tomato^+^ GPC3^+^Hepa1-6 cells, respectively. We found that CAR-macrophages engineered by LNP-GF CAR mRNA strongly inhibited the growth of GPC3^+^Hepa1-6, but not for WT Hepa1-6 cells (Extended Data Fig. 3B). Moreover, we examined the phagocytosis of macrophages by flow cytometry, and found that compared with LNP-Control mRNA, CAR-macrophages engineered by LNP-GF CAR mRNA significantly improved the phagocytosis of GPC3^+^Hepa1-6 cells, but did not improve that of WT Hepa1-6 cells (Figure 4E). Together these data suggested that CAR-macrophages engineered by LNP-GF CAR mRNA could specifically kill tumor cells in an antigen-dependent manner.

**Figure 4:**
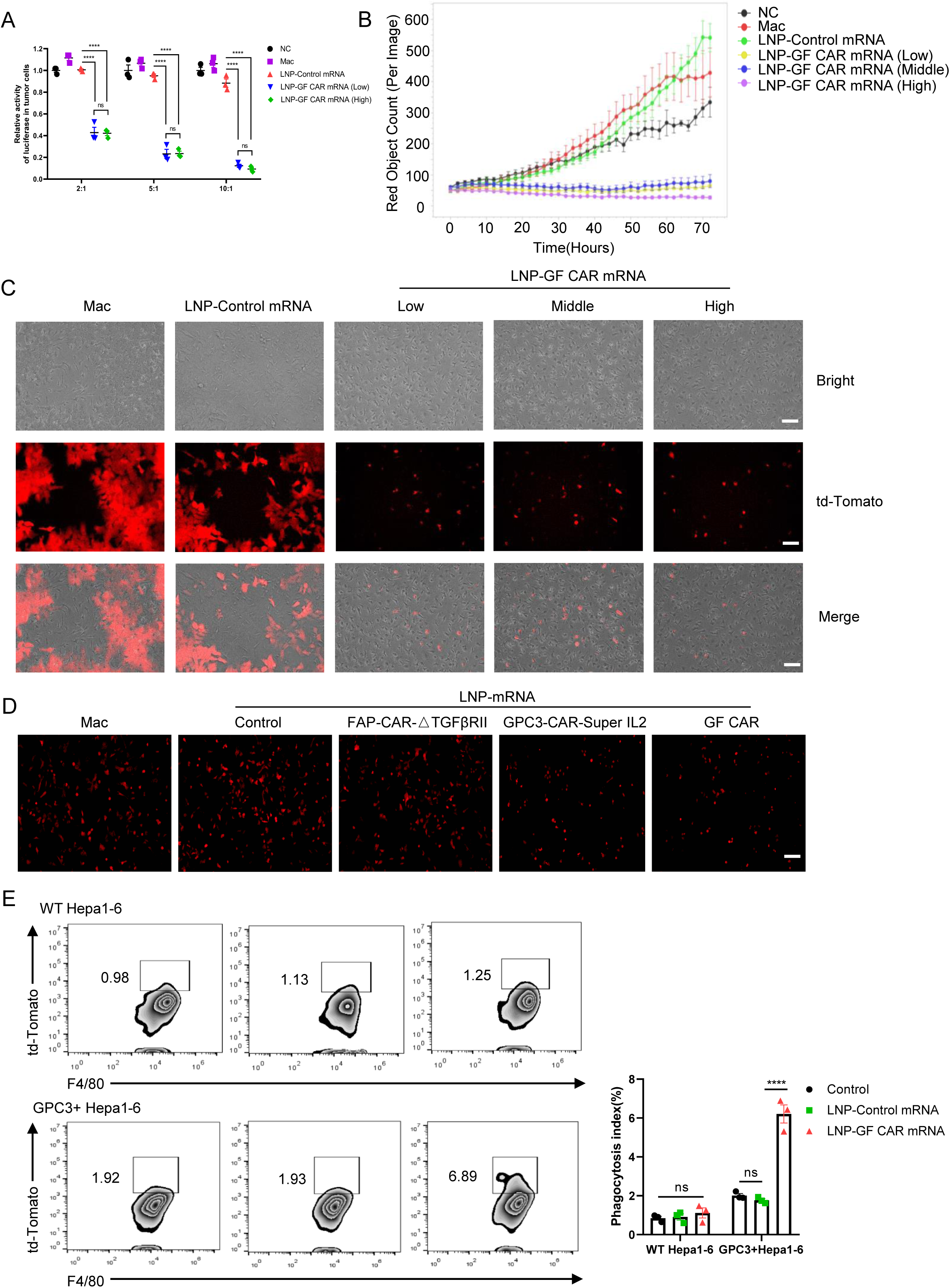
LNP-GF CAR mRNA engineered CAR macrophages specifically and strongly inhibited tumor cell growth. A: BMDMs were transfected with various LNPs for 24 hours. Engineered macrophages were co-incubated with GPC3^+^ luciferase^+^ Hepa1-6 cells at E/T of 2/1, 5/1, and 10/1. After 48 hours of co-incubation, the killing activity of macrophages on tumor cells was analyzed by detecting luciferase activity. The experimental group containing the tumor cells without macrophages was used as an NC. Low dose: 1ug/ml; High dose: 4ug/ml. B: The growth curve of GPC3^+^td-Tomato^+^ Hepa1-6 cells in a time course of ∼3 d after co-incubation in the Incucyte. BMDMs were transfected with various LNPs with td-Tomato^+^ Hepa1-6 cells at an E/T ratio of 10/1 (*n* = 6 replicate wells per group). The Hepa1-6 cells were detected via the td-Tomato signal. Signal capturing was conducted every 2 hours. Low dose: 1ug/ml; Middle dose: 2ug/ml; High dose: 4ug/ml. C: BMDMs were transfected with various LNPs for 24 hours and subsequently co-incubated with GPC3^+^ td-Tomato^+^ Hepa1-6 cells at an E/T ratio of 10/1. Fluorescence microscope imaging was performed to analyze the phagocytosis of macrophages against tumor cells (red represents tumor cell signal). Scale bar: 20 μm D: BMDMs were transfected with LNP-Control mRNA, LNP-FAP-CAR-△TGFβRⅡ mRNA, LNP-GPC3-CAR-Super IL-2 mRNA, and LNP-GF CAR mRNA (1ug/ml) for 24 hours. Fluorescence microscope imaging was performed to analyze the killing activity of CAR-macrophages on GPC3^+^ td-Tomato^+^ Hepa1-6 cells (red represents tumor cell signal). Scale bar: 50 μm E: BMDMs were transfected with LNP-Control mRNA or LNP-GF CAR mRNA (1ug/ml) for 24 hours and subsequently co-incubated with WT Hepa1-6 or GPC3^+^ Hepa1-6. The phagocytosis of macrophages was detected after 48 hours of co-incubation by flow cytometry. Data represent the mean ± s.e.m. from a representative experiment (*n* = 3 biologically independent samples) of two independent experiments. Significance was calculated with two-way ANOVA analysis with multiple comparisons. ns: not significant; \*\*\*\**P* < 0.0001.

### The LNP-GF CAR mRNA achieved complete regression of liver tumor and it depended on the *in vivo* edited macrophages

To investigate whether *in vivo* engineered CAR-macrophages by LNP-GF CAR mRNA can inhibit tumor growth, we established an orthotopic HCC tumor model using Luciferase^+^ GPC3^+^ Hepa1-6 cells. PBS, LNP-Control mRNA, or LNP-GF CAR mRNA were injected intravenously to treat the tumor-bearing mice, and live animal imaging was conducted to monitor the tumor growth (Figure 5A). We found that the LNP-Control mRNA could not inhibit tumor growth, while the LNP-GF CAR mRNA treated group had a remarkable anti-tumor effect, namely, complete remission in 4 out of 5 mice (Figure 5B). Consistent with this result, there were visible tumor sites on the liver of mice in the PBS group and the LNP-Control mRNA group, whereas the liver surface of the LNP-GF CAR mRNA-treated complete responders was smooth, and only one mouse in that group had a small tumor site on the liver (Figure 5C). We also examined whether LNP-GF CAR mRNA had any significant toxic effects on the mice. At the end of the treatment, the weight of mice in the PBS group and the LNP-Control mRNA group did not change significantly. However, compared with the PBS group, the weight of mice in the LNP-GF CAR mRNA group increased significantly, suggesting the weight protective effect of the LNP-GF CAR mRNA (Extended Data Fig. 4A). The liver and renal function of the mice were further analyzed and there were no significant changes in the serum levels of Alanine Aminotransferase (ATL), Aspartate Aminotransferase (AST), Creatinine (Cr), and Urea in all treatment groups (Extended Data Fig. 4B). In addition, HE staining was performed and showed no significant changes in the heart, lungs, spleen, and kidneys among the groups (Extended Data Fig. 4C). These data indicated that LNP-GF CAR mRNA showed complete regression of solid tumors without significant side effects.

**Figure 5:**
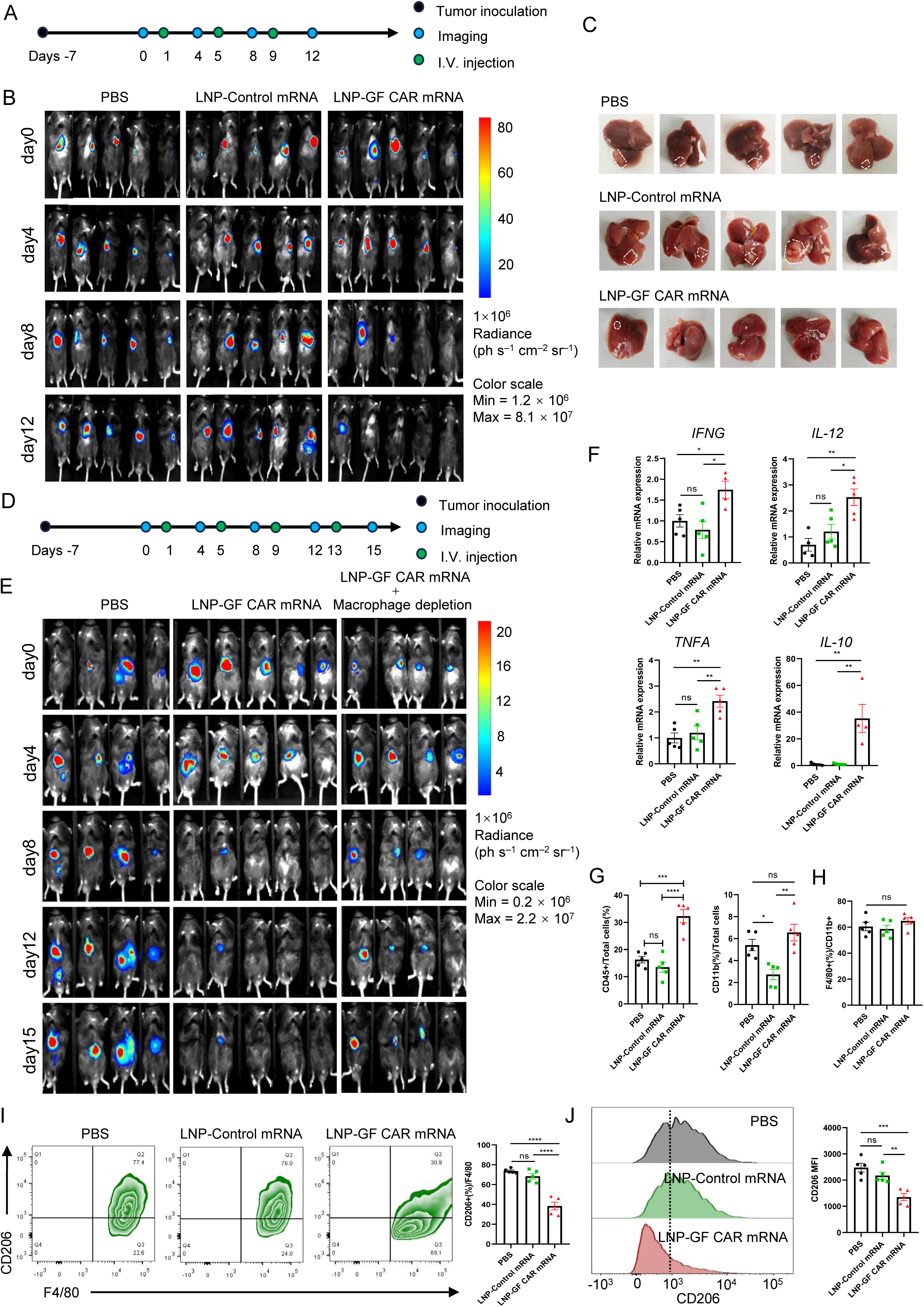
LNP-GF CAR mRNA engineered CAR macrophages *in vivo* eliminate liver tumors. A: An overview of the antitumor activity experiment using LNP-GF CAR mRNA in a GPC3^+^ luciferase^+^ Hepa1-6 orthotopic HCC tumor model. B: Live animal imaging showing the time-dependent change of bioluminescence (BLI) from the orthotopic HCC tumor model after being treated by PBS, LNP-Control mRNA, and LNP-GF CAR mRNA (n=5). C: Photographs of the liver in the orthotopic HCC tumor model in each treatment group, the position circled by the white line indicated the tumor area. D: An overview of the antitumor activity experiment using LNP-GF CAR mRNA under the case of macrophage depletion in a GPC3^+^ Luciferase^+^Hepa1-6 orthotopic HCC tumor model. E: In the case of macrophage depletion, the anti-tumor effect of LNP-GF CAR mRNA was detected by live animal imaging. PBS: (n=4), Macrophage depletion: (n=4), LNP-GF CAR mRNA: (n=5). F: qRT–PCR analysis showing the mRNA expression of cytokines (*IFNG*, *IL-12*, *TNFA*, *IL-10*) in the liver of orthotopic HCC tumor mice in each treatment group (n=5). G: Flow cytometric analysis of the proportion of immune cells (CD45^+^) and myeloid cells (CD45^+^CD11b^+^) in total cells of the liver from the orthotopic HCC tumor model (n=5). H: The proportion of macrophages (CD45^+^CD11b^+^F4/80^+^) in myeloid cells (CD45^+^CD11b^+^), as determined by flow cytometry (n=5). I: Flow cytometric analysis of the proportion of CD206^+^ cells in macrophages of the liver from the orthotopic HCC tumor model (n=5). J: Expression levels of CD206 on macrophages of liver from the orthotopic HCC tumor model, as determined by flow cytometry (n=5). Significance was calculated with one-way ANOVA analysis with multiple comparisons and is presented as mean ± s.e.m., ns: not significant; *p < 0.05, **p < 0.01, ***p < 0.001, ****p < 0.0001.

Next, we explored whether the anti-tumor effect of LNP-GF CAR mRNA was dependent on macrophages. To determine whether intravenous LNP mainly edits liver macrophages, we mixed a certain proportion of DiD dye into LNPs, so that the cells in the liver that ingested LNPs would be labeled by DiD. 24 hours after intravenous LNP injection, the liver cells were analyzed by flow cytometry. The result showed that macrophages accounted for 70% of all DiD-positive cells (Extended Data Fig. 5A), indicating that intravenous LNPs are indeed mainly taken up by macrophages in the liver and that LNP-GF CAR mRNA mainly edited liver macrophages. To determine whether the anti-tumor effect of LNP-GF CAR mRNA was primarily dependent on the *in vivo* edited CAR-macrophages, we performed the anti-tumor animal experiment again after depletion of macrophages by administration of Clodronate Liposomes[26] (Figure 5D). As the previous results, LNP-GF CAR mRNA significantly inhibited tumor growth, and four out of the five mice achieved complete cure. However, in the case of macrophage depletion, LNP-GF CAR mRNA only showed a weaker effect in repressing the tumor (Figure 5E), demonstrating that the anti-tumor effect of LNP-GF CAR mRNA was mainly dependent on the production of endogenous CAR-macrophages.

### LNP-GF CAR mRNA promoted immune activation and inhibits M2 polarization of macrophages

Next, we analyzed the effect of LNP-GF CAR mRNA on the immune function of mice. Interestingly, we observed that the spleens in the LNP-GF CAR mRNA treated group were significantly enlarged and their weight significantly increased compared with those in the PBS and LNP-Control mRNA groups (Extended Data Fig. 6A, B), suggesting that LNP-GF CAR mRNA might have stimulated the immune system of the mice. In addition, we analyzed the expression of inflammatory factors in liver tissues and found that LNP-GF CAR mRNA significantly increased the mRNA expression of *IFNG*, *IL-12*, *TNFA*, and *IL-10* (Figure 5F). These results indicated that the immune system of the LNP-GF CAR mRNA-treated group was significantly activated.

Thus we further explored the effect of LNP-GF CAR mRNA on the immune cell population in the liver. The results showed that compared with the PBS group and the LNP-Control mRNA group, the LNP-GF CAR mRNA treated group significantly increased the proportion of the CD45 immune cell population but there was no significant effect on myeloid cell population (Figure 5G), suggesting that LNP-GF CAR mRNA altered the number of global immune cells in the mouse liver, but it was not caused by the myeloid cells. Next, we assessed the polarization changes of CAR-macrophages engineered by LNP-GF CAR mRNA *in vivo*. The results showed that even though the proportion of macrophages to myeloid cells did not change among the groups (Figure 5H), the proportion of CD206^+^ macrophages in the LNP-GF CAR mRNA treatment group was markedly reduced, and the average fluorescence intensity of CD206 on the surface of macrophages was also significantly decreased, compared with the control groups (Figure 5I, J), suggesting that the LNP-GF CAR mRNA treatment could robustly inhibit the M2 polarization of macrophages. This might also be a result by the obtained tolerance against the TGFβ signaling of the CAR-macrophages engineered by LNP-GF CAR mRNA *in vivo*.

### *In vivo* engineered CAR-macrophages broke the physical barrier of solid tumors

We next delved into the anti-tumor mechanisms of *in vivo* engineered CAR-macrophages. Due to the FAP-CAR and △TGFβRII design, we focused on analyzing whether CAR-macrophages engineered by LNP-GF CAR mRNA *in vivo* could reduce the physical barrier constructed by CAFs outside the tumor. Because TGFβ is known to activate CAFs and promote M2 polarization of macrophages, we first determined whether CAR-macrophages engineered by LNP-GF CAR mRNA had decreased TGFβ signaling. Immunofluorescence staining of liver tissues for F4/80 and p-SMAD2/3 showed that the signal of p-SMAD2/3 in macrophages in the LNP-GF CAR mRNA treated group was significantly reduced (Figure 6A). In addition, we were surprised to find that TGFβ signaling in other cells besides macrophages was also considerably reduced in the LNP-GF CAR mRNA treated group. To validate this phenomenon, we performed p-SMAD2/3 immunohistochemistry on the liver to analyze the overall TGFβ signaling again. The results also showed that the LNP-GF CAR mRNA treated group significantly reduced p-SMAD2/3 at the junction of tumor and healthy tissue (Figure 6B). This indicated that LNP-GF CAR mRNA engineered CAR-macrophages not only tolerated the TGFβ signal themselves but also led to the inhibition of the TGFβ signal in the overall liver tissue. Next, we focused on whether the CAFs layer surrounding the tumor was disrupted. By Sirius red staining of liver tissues, we found that the fibrotic area was significantly reduced in the LNP-GF CAR mRNA treated group, and the thickness of the peripheral fibrocyte band was much thinner (Figure 6C, D). To prove the specific elimination of FAP-positive cells, we examined the expression of FAP in liver tissue and found that LNP-GF CAR mRNA effectively reduced the expression level of FAP (Figure 6E). Besides, with immunofluorescence staining of FAP, we observed that the FAP-positive region was markedly reduced in the LNP-GF CAR mRNA treated group, and there were almost no cell bands composed of FAP-positive CAFs in the tumor periphery in the treated group (Figure 6F, G). Combining the above data, we demonstrated that LNP-GF CAR mRNA engineered CAR-macrophages broke down the physical barrier constructed by CAFs cells outside the tumor. The removal of the physical barrier could allow infiltration of CD8^+^ T cells into the tumor. Therefore, we analyzed the distribution of CD8^+^ T cells, and the results showed that a large number of CD8^+^ T cells in the PBS and the LNP-Control mRNA groups were retained in the FAP-positive cell region, whereas the LNP-GF CAR mRNA treated group showed markedly increased infiltration of CD8^+^ T cells into the tumor (Figure 6H). Results showed that LNP-GF CAR mRNA treated group significantly increased the number of CD8^+^ T cells in the tumor core area (Figure 6I). The above data illustrated that CAR-macrophages engineered by LNP-GF CAR mRNA could break the physical barrier constructed by FAP-positive CAFs, reduce the TGFβ signal in the liver, and rescue CD8^+^ T cells from the stroma region. These findings laid the foundation for the reactivation of CD8^+^ T cells and the unleash of their long lasting anti-tumor immunity.

**Figure 6:**
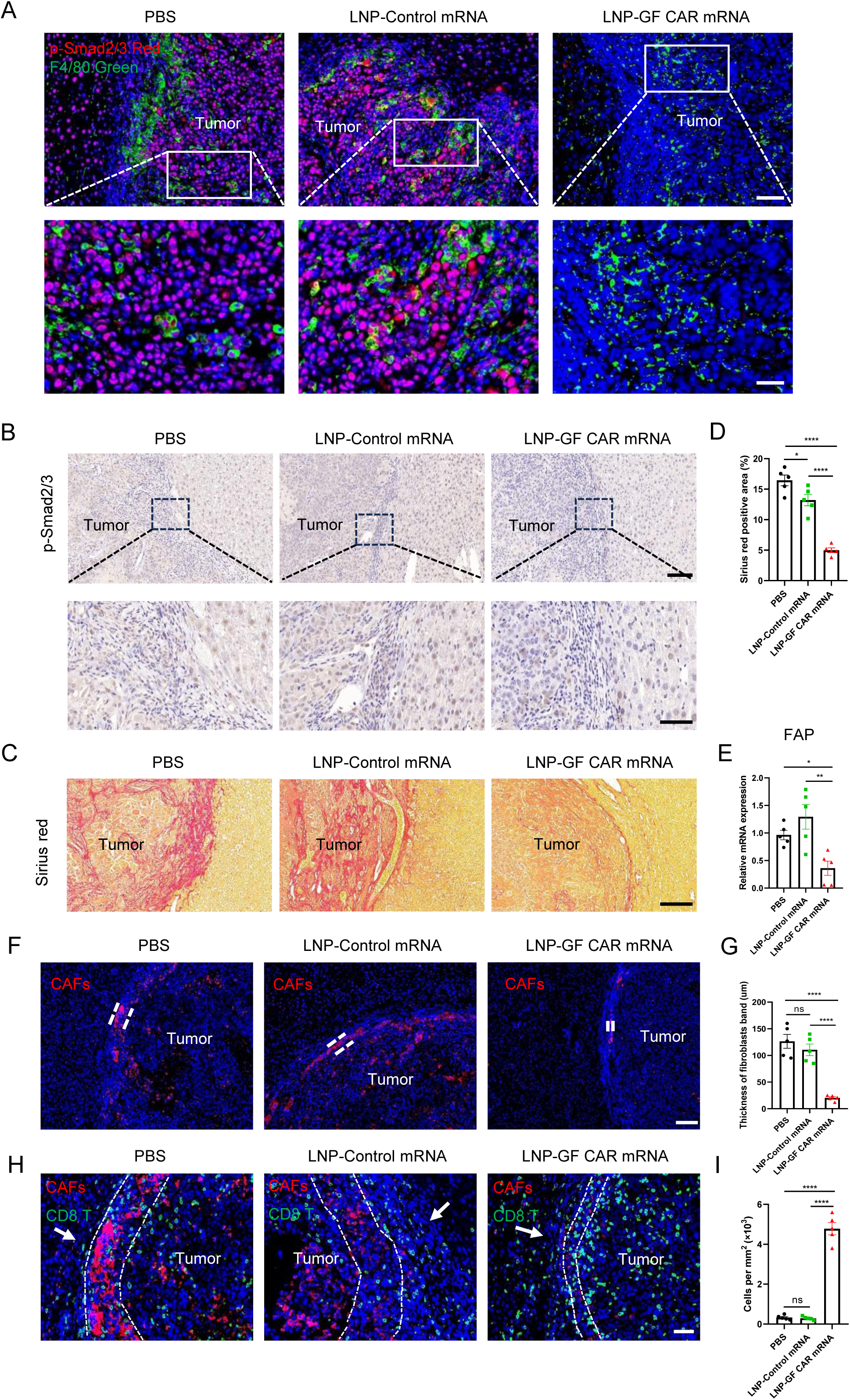
*In vivo* engineered CAR macrophages by LNP-GF CAR mRNA break down the physical barrier of solid tumors. A: Immunofluorescence staining of F4/80, p-SMAD2/3 in the liver tissue sections of orthotopic HCC tumor mice. Scale bar: 50 μm, magnified partial view Scale bar: 20 μm. B: Immunohistochemical staining of p-SMAD2/3 in the liver tissue sections of orthotopic HCC tumor mice. Scale bar: 100 μm, magnified partial view Scale bar: 50 μm. C, D: Sirian red staining of the liver in each treatment group of orthotopic HCC tumor mice and quantification of the Sirius red-positive area. Scale bar: 100 μm E: qRT–PCR analysis showing the mRNA expression of FAP in the liver of orthotopic HCC tumor mice (n=5). F, G: Immunofluorescence staining of FAP (Red) in the liver of orthotopic HCC tumor mice and quantification of the thickness of CAFs layer. Scale bar: 100 μm H, I: The distribution of CD8^+^ T cells in the liver of orthotopic HCC tumor mice was analyzed by immunofluorescence assay (H) and quantification of the CD8^+^ T cells in the tumor core area (I). CD8 (Green), FAP (Red). The white arrow indicates the direction of CD8^+^ T infiltration and the white dotted line indicates the peripheral area of the tumor. Scale bar: 50 μm. Significance was calculated with one-way ANOVA analysis with multiple comparisons and is presented as mean ± s.e.m., ns: not significant; *p < 0.05, **p < 0.01, ***p < 0.001, ****p < 0.0001.

### Transient-engineered CAR-macrophages *in vivo* are sufficient to form immune memory

Moreover, according to our design, the *in vivo* engineered CAR-macrophages were able to secrete Super IL-2 *in situ*, so we further analyzed the changes of the T cell populations. Further analyzed by flow cytometry, we found that LNP-GF CAR mRNA significantly increased the proportion of total CD3 T cells in the liver (Figure 7A). Combined with the previous data, this suggested that the increase of global immune cells induced by the LNP-GF CAR mRNA treatment was mainly contributed by the increase in the proportion of T cells. Further analysis showed that LNP-GF CAR mRNA significantly increased the proportion of CD4^+^, CD8^+^, and IFNγ^+^CD8^+^ cells (Figure 7B-D). In contrast, LNP-GF CAR mRNA significantly decreased the proportion of Treg cells in CD4^+^ T cells compared with LNP-Control mRNA (Figure 7E). These results suggested that LNP-GF CAR mRNA had an effect on preferentially activating CD8^+^ T cells. In summary, these results demonstrated that LNP-GF CAR mRNA could not only promote the proliferation and infiltration of CD8^+^ T cells but also reactivate the immune function of CD8^+^ T cells.

**Figure 7:**
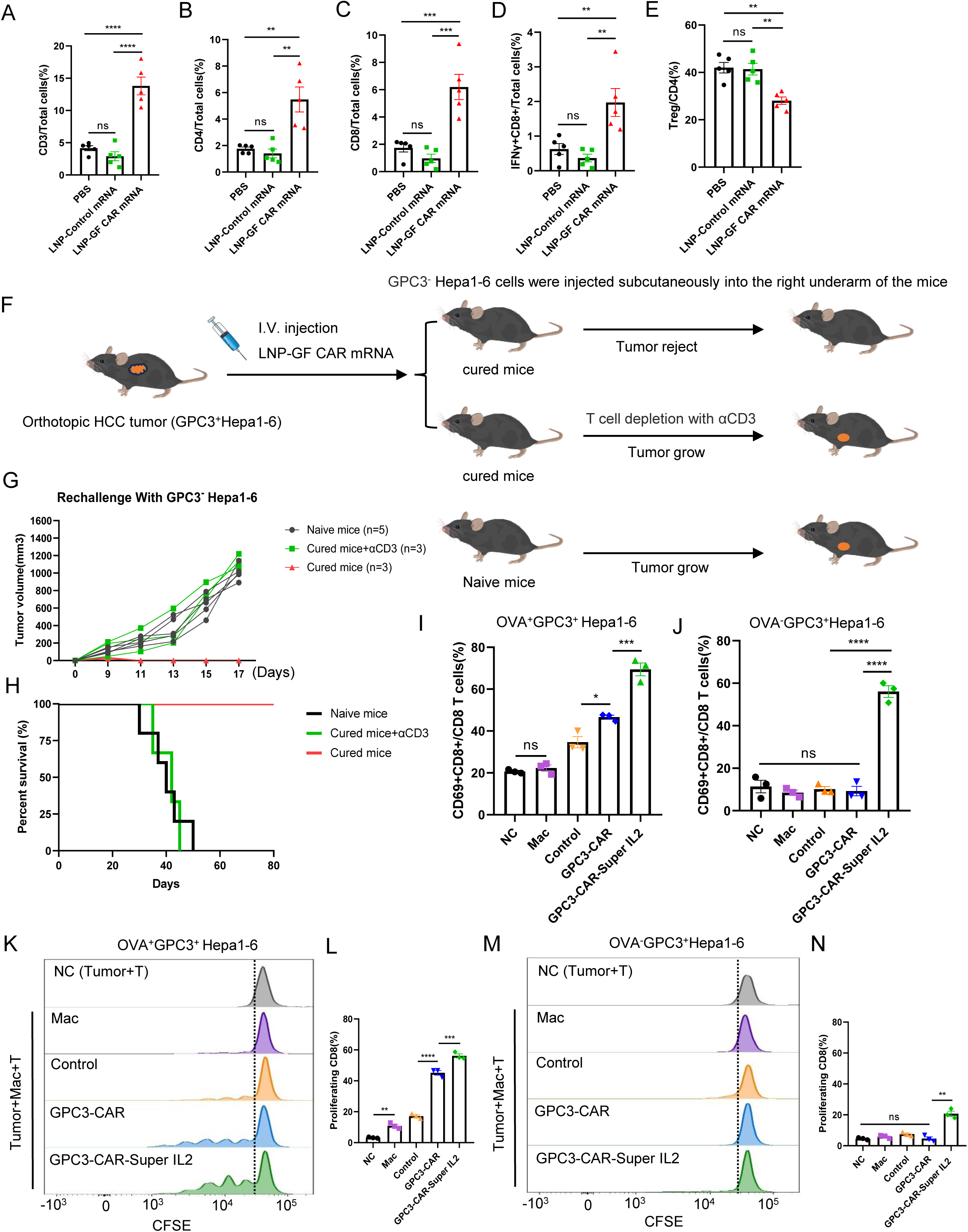
*In vivo* engineered CAR macrophages preferentially activate CD8^+^ T cells and promote the formation of T cell immune memory. A-D: Flow cytometric analysis of the proportion of CD3^+^ T cells, CD4^+^ T cells, CD8^+^ T cells, and IFNγ^+^ CD8^+^ T cells in total cells of liver from the orthotopic HCC tumor model (n=5). E: The proportion of Treg cells in CD4^+^ T cells of the liver from the orthotopic HCC tumor model was analyzed by flow cytometry (n=5). F: Schematic of rechallenge experiments using fully responder mice cured by LNP-GF CAR mRNA in the orthotopic HCC tumor model. The rechallenge was injected subcutaneously using GPC3^−^Hepa1-6 (GPC3 negative) cells one month post complete remission. G: 1×10^6^ GPC3^−^Hepa1-6 cells were injected subcutaneously in cured and naive mice. The volume of subcutaneous tumors in each group of mice was measured starting on day nine and recorded every two days. H: Kaplan–Meier survival curve of the rechallenge experiments. I-N: BMDMs were transfected with various LNPs for 24 hours and subsequently co-incubated with OVA^+^GPC3^+^Hepa1-6 or OVA^−^GPC3^+^Hepa1-6 cells and T cells derived from OT1 mice. The activation of CD8^+^ T cells (I, J) and proliferation of CD8^+^ T cells indicated by CFSE staining (K, M) were detected after 72 hours of co-incubation by flow cytometry. The quantification of CD8^+^ T cells proliferation was shown (L, N). NC: No macrophages, Mac: Untransfected macrophages, Control: LNP-Control mRNA transfected macrophages, GPC3-CAR: LNP-GPC3-CAR mRNA transfected macrophages, GPC3-CAR-Super IL-2: LNP-GPC3-CAR-Super IL-2 mRNA transfected macrophages. Significance was calculated with one-way ANOVA analysis with multiple comparisons and is presented as mean ± s.e.m., ns: not significant; *p < 0.05, **p < 0.01, ***p < 0.001, ****p < 0.0001.

The formation of immune memory is essential to prevent tumor recurrence and metastasis. In addition, we have demonstrated that LNP-GF CAR mRNA confers CAR-macrophages with the ability to promote CD8^+^ T cells proliferation and activation. To further investigate whether *in vivo* engineered CAR-macrophages could promote T cell immune memory formation, we performed a rechallenge assay in cured mice by subcutaneous injection of GPC3^−^Hepa1-6 cells (Figure 7F). We found that cured mice in the orthotopic HCC tumor model completely rejected the growth of subcutaneous GPC3^−^Hepa1-6 tumors (Figure 7G). However, in the case of T cell depletion, the cured mice resumed the growth of subcutaneous tumors and the tumor growth rates were similar to those in the naive mice (Figure 7G). Correspondingly, mice cured by LNP-GF CAR mRNA had 100% survival, while the cured mice treated with αCD3 or naive mice all died within two months (Figure 7H). This result strongly demonstrated that mice cured by LNP-GF CAR mRNA in the orthotopic HCC tumor model form T cell immune memory, and the mRNA transfected CAR-macrophages functioned as a tumor vaccine to prevent recurrence. Notably, as GPC3 antigen-negative tumor cells were inoculated in the rechallenge assay, it suggests a potential mechanism of antigen spreading for the activated T cells.

To further elucidate the antigen spreading mechanism of CAR-macrophages, we constructed a Hepa1-6 cell line overexpressing full-length GPC3 and OVA (Ovalbumin) antigens. In addition, we constructed GPC3-CAR molecules without the Super IL-2 sequence to distinguish the contribution of GPC3-CAR versus IL-2 to T cell activation and antigen spreading. We prepared LNPs containing these different mRNA sequences. CAR-macrophages engineered by different LNPs were co-incubated with OVA^+^GPC3^+^Hepa1-6 cancer cells and T cells derived from OT1 mice. After 72 hours of co-incubation, the proliferation and activation of CD8^+^ T and CD4^+^ T cells were detected by flow cytometry. We found that CAR-macrophages engineered by LNP-GPC3 CAR mRNA significantly enhanced both the proliferation and activation of CD8^+^ T cells compared with LNP-Control mRNA (Figure 7I, K, L). CAR-macrophages engineered by LNP-GPC3-CAR-Super IL-2 mRNA further promoted the proliferation and activation of CD8^+^ T cells (Figure 7I, K, L). In contrast, when co-incubated with OVA^−^GPC3^+^Hepa1-6 cells, CAR-macrophages engineered by LNP-GPC3 CAR mRNA did not promote proliferation and activation of CD8^+^ T cells. Only the LNP-GPC3-Super IL-2 CAR mRNA treatment increased the proliferation and activation of CD8^+^ T cells (Figure 7J, M, N). This suggests a general role of Super IL-2 in promoting the activation and proliferation of CD8^+^ T cells. Taken together, these results suggest that the GPC3 targeting CAR-macrophages indeed promoted proliferation and activation of CD8^+^ T cells derived from OT1 mice by presenting the OVA antigen through antigen spreading. In addition, we also observed that macrophages in all treatment groups did not promote the proliferation of CD4^+^ T cells (Extended Data Fig. 7A-D). Since CD4^+^ T cells from OT1 mice do not recognize OVA antigens, this result rigorously demonstrates that the mechanism by which CAR-macrophages activate T cells is through antigen spreading to OVA and its presentation to CD8^+^ T cells.

## Discussion

At present, the success of CAR-T therapy in treating hematopoietic malignancies has been difficult to replicate in solid tumors[27]. The promising CAR-macrophages have also been hindered by lower editing efficiency and high cost of manufacturing. Here, we engineered CAR-macrophages *in vivo* by simultaneously delivering mRNA of GPC3-CAR-Super IL-2 and FAP-CAR-△TGFβRII through the LNP-mRNA system to treat solid tumors. *In vivo* engineered CAR-macrophages demonstrated strong anti-tumor capabilities, achieving an 80% cure rate in the orthotopic HCC tumor model. We observed that CAR-macrophages engineered *in vivo* could break down the physical barrier surrounding solid tumors and promote the infiltration and activation of CD8^+^ T cells. Importantly, we elucidated the association between *in vivo* engineered CAR-macrophages and T cell immune memory, providing new possibilities for preventing tumor recurrence and metastasis.

Increasing evidence demonstrates that CAFs play an important role in the development of solid tumors. CAFs can not only secrete a variety of growth factors to directly nourish the growth of tumor cells, but also cause tolerance to chemotherapy, radiotherapy, and immunotherapy by remodeling ECM[28]. FAP-CAR T cells targeting CAFs have been used to treat solid tumors[29–31]. However, FAP-CAR T cells still had the disadvantages of traditional CAR-T cells, such as T cell exhaustion due to the immunosuppressive tumor microenvironment. In addition, it has been reported that FAP-CAR T cells could cause severe bone marrow cytopenia and cachexia due to their off-target effects[32, 33]. Moreover, we found that the number of endogenous CD8^+^ T cells was very limited and they were mainly confined to the stroma region of the tumor. Spatial transcriptome data of hepatocellular carcinoma patients showed that limited NK cells were also confined to the periphery of the tumor by fibrocyte bands[21]. Therefore, engineered CAR-T cells or CAR-NK cells *in vivo* might not solve the problem of breaking down the physical barrier and infiltrating into solid tumors. However, we showed endogenous macrophages are widely distributed both inside and outside the tumor. Moreover, many studies have shown the interaction between TAMs and CAFs in the process of tumor development[34–36], such as the spatial interaction of FAP^+^ fibroblasts and SPP1^+^ TAMs in patients with colorectal cancer[37]. Therefore, *in vivo* engineered CAR-macrophages may have superior accessibility to break down the physical barrier constructed by CAFs.

TGFβ is the central cytokine in the activation of CAFs and can also inhibit the immune function of macrophages. So we designed a FAP-CAR structure containing △TGFβRII. As expected, we observed that *in vivo* engineered CAR-macrophages by LNP-GF CAR mRNA significantly reduced p-SMAD2/3 signaling. Moreover, the overall level of p-SMAD2/3 in the liver of the LNP-GF CAR mRNA treatment group was also significantly reduced. It might be due to the uptake of LNP-GF CAR mRNA by different types of cells in the liver. However, our data showed that 70% of the cells ingesting LNPs were macrophages. Therefore, we suspect that the abundant macrophage population competitively binds TGFβ by overexpressing △TGFβRII, thereby reducing TGFβ’s availability to other cells, which can be considered as the ‘sponge effect’.

mRNA is expressed in the cytoplasm without the risk of genome insertion, which brings safety but also has the problem of short expression time[38]. Therefore, *in vivo* engineered CAR-macrophages by LNP-GF CAR mRNA were transient and would gradually fade away. This has raised concerns about the recurrence and metastasis of tumors. However, our study demonstrated that transiently engineered CAR-macrophages *in vivo* were sufficient to form a long-lasting T cell immune memory. It was worth noting that *in vivo* engineered CAR-macrophages by LNP-GF CAR mRNA could secrete Super IL-2 *in situ.* In addition to promoting the proliferation and activation of T cells, IL-2 also contributes to the formation of memory CD8^+^ T cells[39]. IL-2 could reduce the apoptosis of antigen-specific CD8^+^ T cells and increase the number of memory CD8^+^ T cells during the regression phase of the immune response[40]. Most importantly, CAR-macrophages could spatially bring tumor cells and T cells close to each other and activate T cells through antigen presentation. The spatial proximity of these cells, combined with *in-situ* Super IL-2 secretion, may synergistically contribute to the formation of memory CD8^+^ T cells.

Solid tumors are highly heterogeneous in terms of the types of antigen each tumor cell expresses. A single targeting or even dual targeting strategy with conventional CAR-T therapies could not solve this challenge. Our *in vivo* engineered CAR-macrophage platform provides a solution to accomplish both antigen-specific killing and unspecific tumor cell killing, and the latter is conferred by the capacity of macrophages to phagocytose tumor cells and present extra tumor antigens to stimulate a more diverse pool of T cells, a process called antigen spreading. Ultimately, the tumor cells that do not carry the original CAR-targeting antigen can also be eliminated. Thus, this platform is a showcase of the route toward solving solid tumor heterogeneity by those targeted immune cell therapy approaches.

## Acknowledgements

We thank Na Kong (Zhejiang University) for providing the protocol of LNP preparation, Shuai Liu (Zhejiang University) and Xin Lin (Tsinghua University) for discussion. This work was funded by the National Natural Science Foundation of China (82373238, 31871453, and 91857116 to J.Z.), the National Key Research and Development Program of China (2018YFA0107103 and 2018YFC1005002 to J.Z.), Zhejiang Innovation Team grant (2019R01004 to J.Z.), the Zhejiang Natural Science Foundation (LR19C120001 to J.Z.), the Key Research and Development Program of Zhejiang Province (2023C03036).

## Author contributions

J.Z. and S.Z. designed the study and drafted the manuscript. S.Z. conducted the majority of the experiments. H.L, H.Z participated in animal experiments and constructed tumor cell lines. X.D constructed cell lines of overexpressed OVA and preparation of OT1 mice. Q.W participated in animal experiments. A.L. participated in research discussions.

## Methods

### Mice and cell culture

Five-to six-week-old male C57BL/6 mice were purchased from Zhejiang Vital River Laboratory Animal Technology Co., Ltd. T cell antigen receptor-transgenic OT-I mice (C57BL/6-Tg(TcraTcrb)1100Mjb/J) were purchased from Jackson Laboratory. The Hepa1-6 and 3T3 cell lines were purchased from ATCC. Cells were maintained in complete DMEM with 10% fetal bovine serum (Homeland, China), 200 mM L-glutamine, and 100 units/ml penicillin-streptomycin). All cells used in this study were cultured at 37 °C with 5% CO2.

### Generation of stable tumor cell lines

The GPC3 coding sequence from human HepG2 cells was cloned into the Lenti-EF1α-PGK-Puro vector and overexpressed in Hepa1-6 cells using the abovementioned strategy to establish the GPC3^+^Hepa1-6 cell line. 3T3 cell lines overexpressing mouse FAP-GFP were prepared by the same method.

### Macrophage distribution experiment *in vivo*

BMDMs were stained with DiD dye. Then BMDMs were collected after being washed three times with PBS. For mice with established subcutaneous tumor models or orthotopic HCC tumor models, DiD-labeled macrophages were injected intravenously. The tissues (Heart, liver, spleen, lung, kidney, and brain) of mice were imaged 3 days later.

### Western blotting

Western blotting was performed according to the standard protocol. The primary antibodies and dilutions used were as follows: p-SMAD2/3 (1:1000), GAPDH (1:3000).

### Immunohistochemistry, and Immunofluorescence staining

Tissues were harvested, extensively washed with PBS, and immediately immersed in 4% paraformaldehyde. The fixed tissue was sent to Shanghai Ruiyu Biotechnology Co., Ltd. (Shanghai, China), for immunohistochemistry and immunofluorescence staining. The CD31 staining represents blood vessels. DiD signal (autofluorescence) represents the location of foreign macrophages. F4/80 and CD8 staining represents the distribution of endogenous macrophages and CD8 T cells. FAP staining represents the distribution of CAFs. p-SAMD2/3 staining represents the activation of the TGFβ signal. The Sirius red stain represents fibrotic areas.

### RNA isolation and qRT–PCR

Total RNA extraction was performed using Eastep Super Total RNA Extraction Kit (Shanghai Promega Bio, LS1040) according to the manufacturer’s instructions. cDNA was prepared using HiScript II Q Select RT SuperMix for qRT–PCR with gDNA wiper (Vazyme, R223-01). Gene expression was analyzed in triplicates using HiScript II One Step qRT– PCR SYBR Green Kit (Vazyme, Q221-04) and Bio-Rad PCR machine (CFX-96 Touch).

## Figure Legend

**Supplementary Figure 1:**
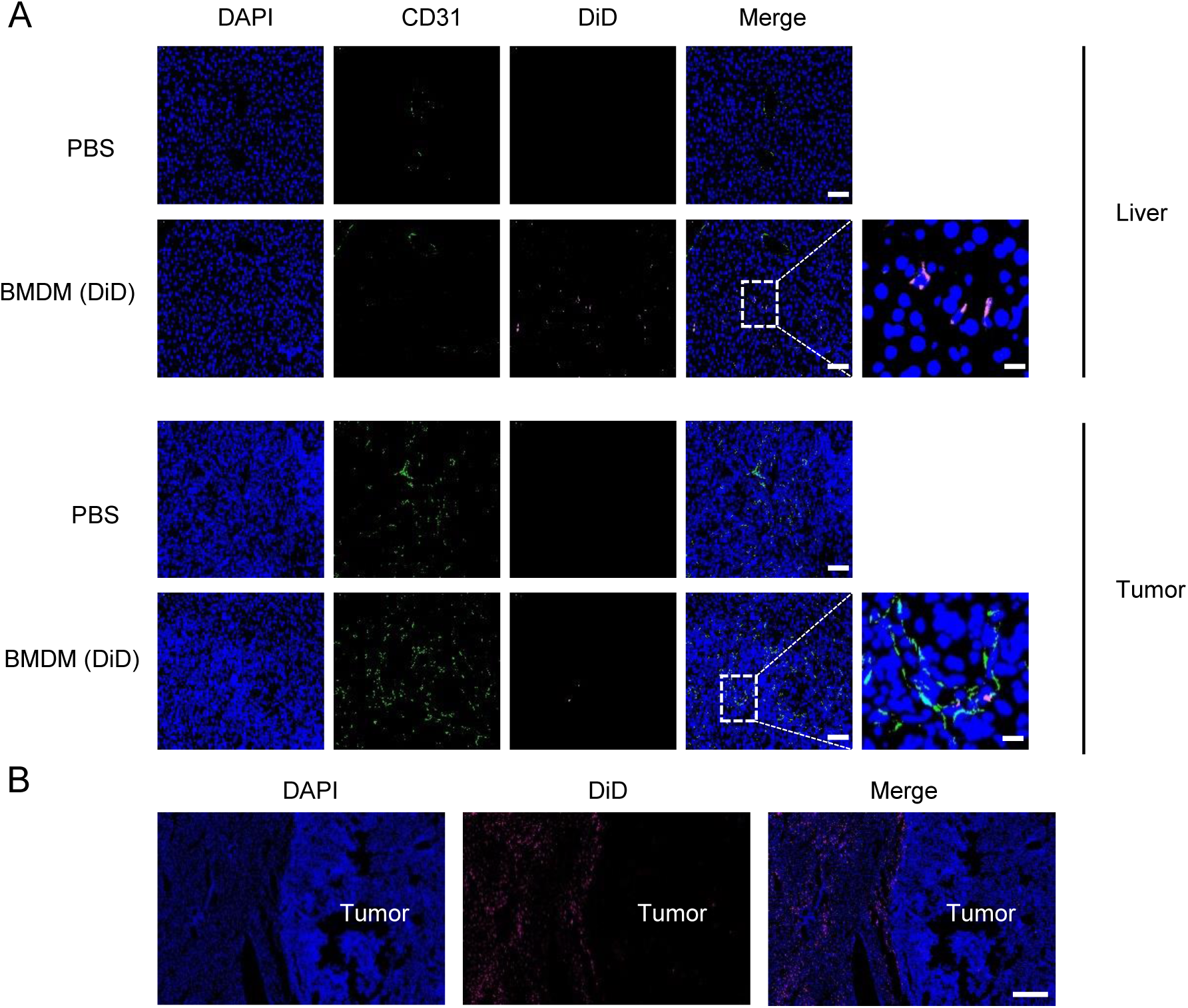
Distribution of exogenous macrophages in solid tumors. A: Immunofluorescence assays showing the distribution of DiD-labeled BMDMs in the liver and tumor. CD31 (Green) indicated blood vessels and DiD (pink) indicated BMDMs. Scale bar: 50 μm, magnified partial view Scale bar: 10 μm. B: Immunofluorescence assays showing the distribution of DiD-labeled BMDMs in the liver of orthotopic HCC tumors. DiD signal (pink) indicated the location of BMDMs. Scale bar: 300 μm.

**Supplementary Figure 2:**
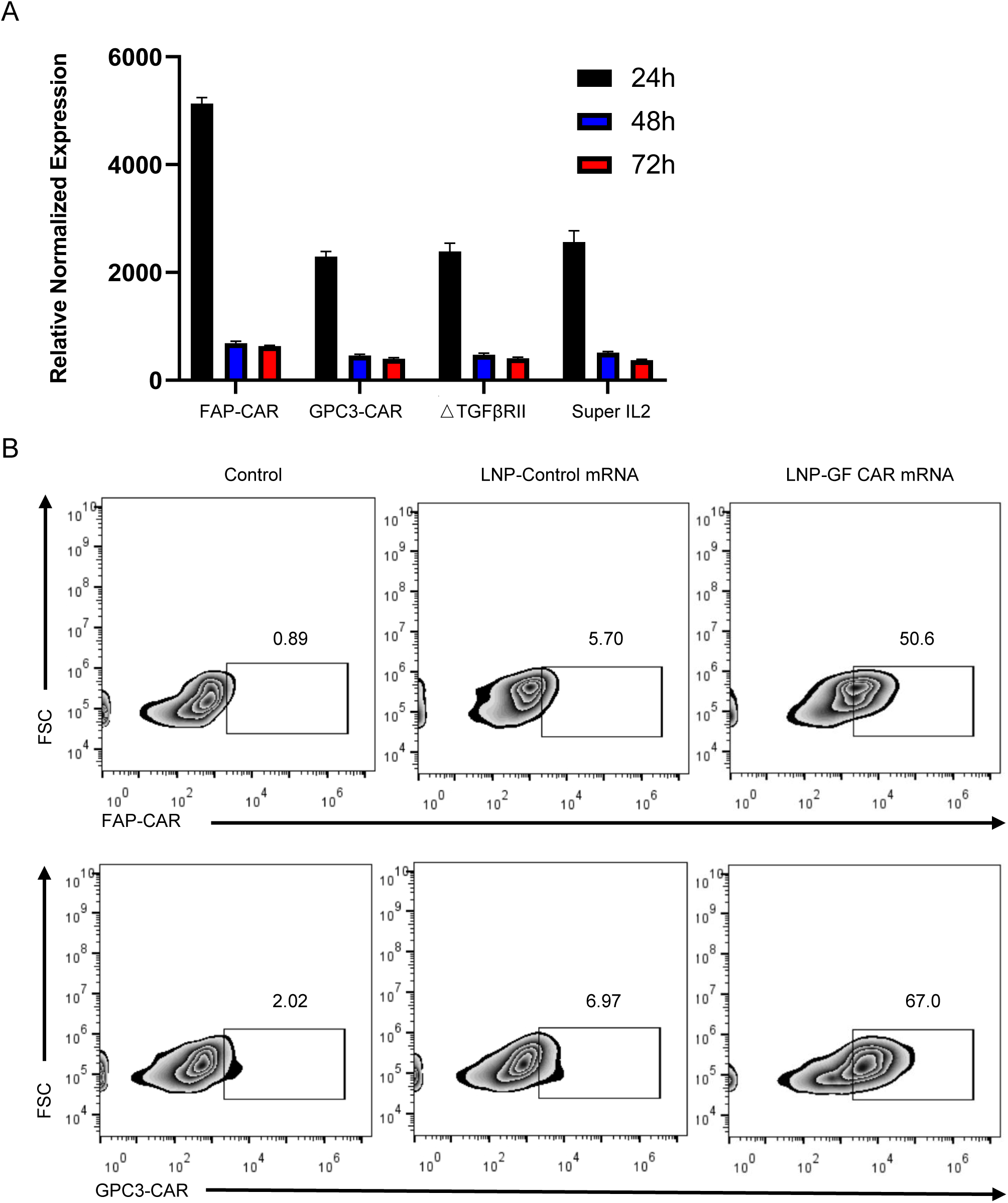
Editing efficiency of CAR macrophages by LNP-GF CAR mRNA. A: BMDMs were transfected with LNP-GF CAR mRNA(1ug/ml) for 24 hours, the mRNA expression of FAP-scfv, GPC-scfv, △TGFβRⅡ, and Super IL-2 were detected by qRT-PCR at different time points. B: BMDMs were transfected with LNP-GF CAR mRNA(1ug/ml) for 24 hours. The expression of GPC3-CAR and FAP-CAR on macrophage surface was analyzed by flow cytometry.

**Supplementary Figure 3:**
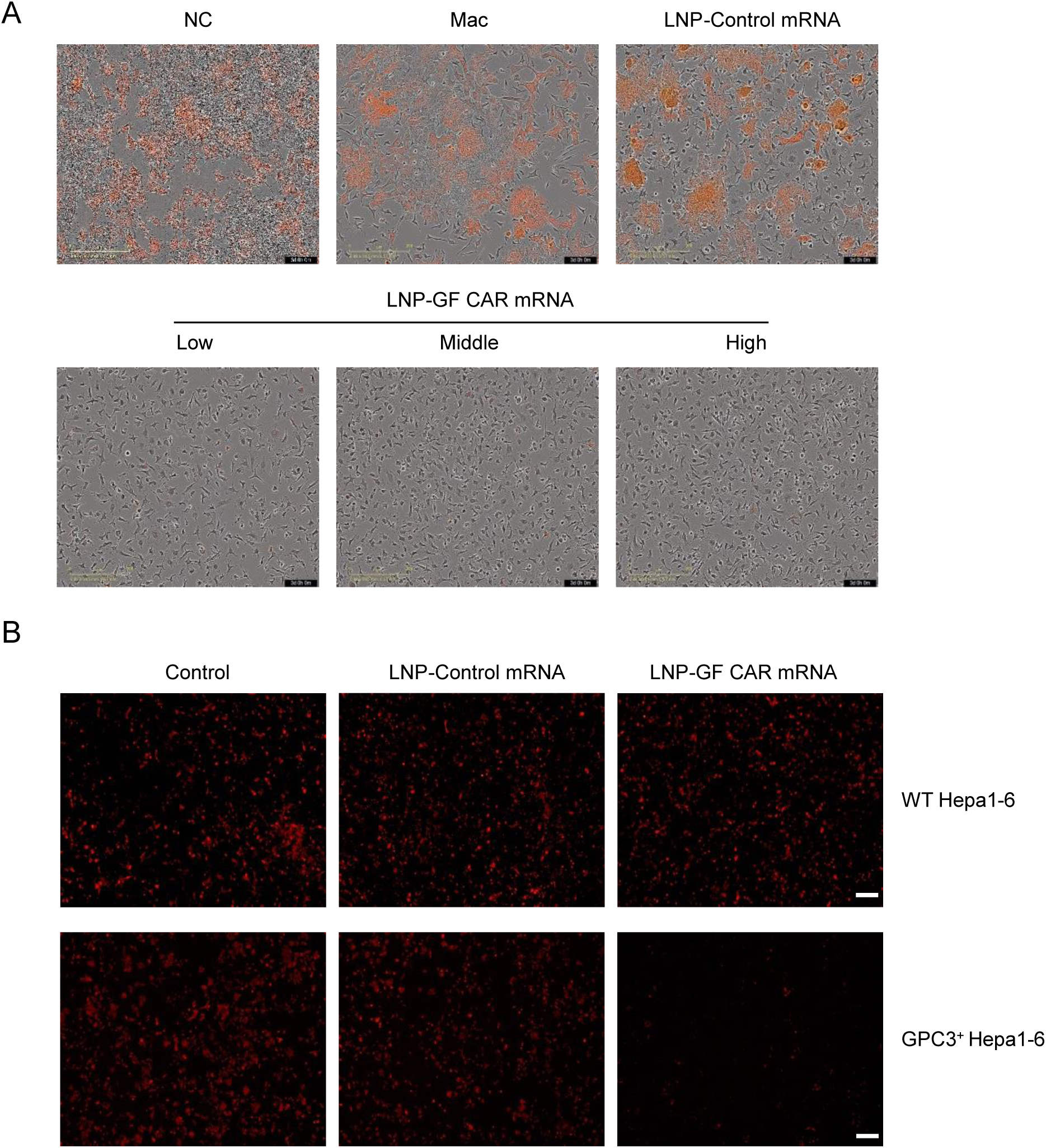
CAR macrophages engineered by LNP-GF CAR mRNA destroy tumor cells in an antigen-dependent manner. A: BMDMs were transfected with various LNPs for 24 hours and subsequently co-incubated with GPC3^+^ td-Tomato^+^ Hepa1-6 cells at an E/T ratio of 10/1. At the 72-hour coculture endpoint, Incucyte imaging data on each group of cells were shown (red signal represents tumor cells). B: BMDMs were transfected with various LNPs for 24 hours and subsequently co-incubated with GPC3^+^ td-Tomato^+^ Hepa1-6 cells or WT td-Tomato^+^Hepa1-6 cells at an E/T ratio of 10/1. After 48 hours, fluorescence microscopy was used to analyze the killing effect of macrophages on tumor cells (red signal represents tumor cells). Scale bar: 100 μm.

**Supplementary Figure 4:**
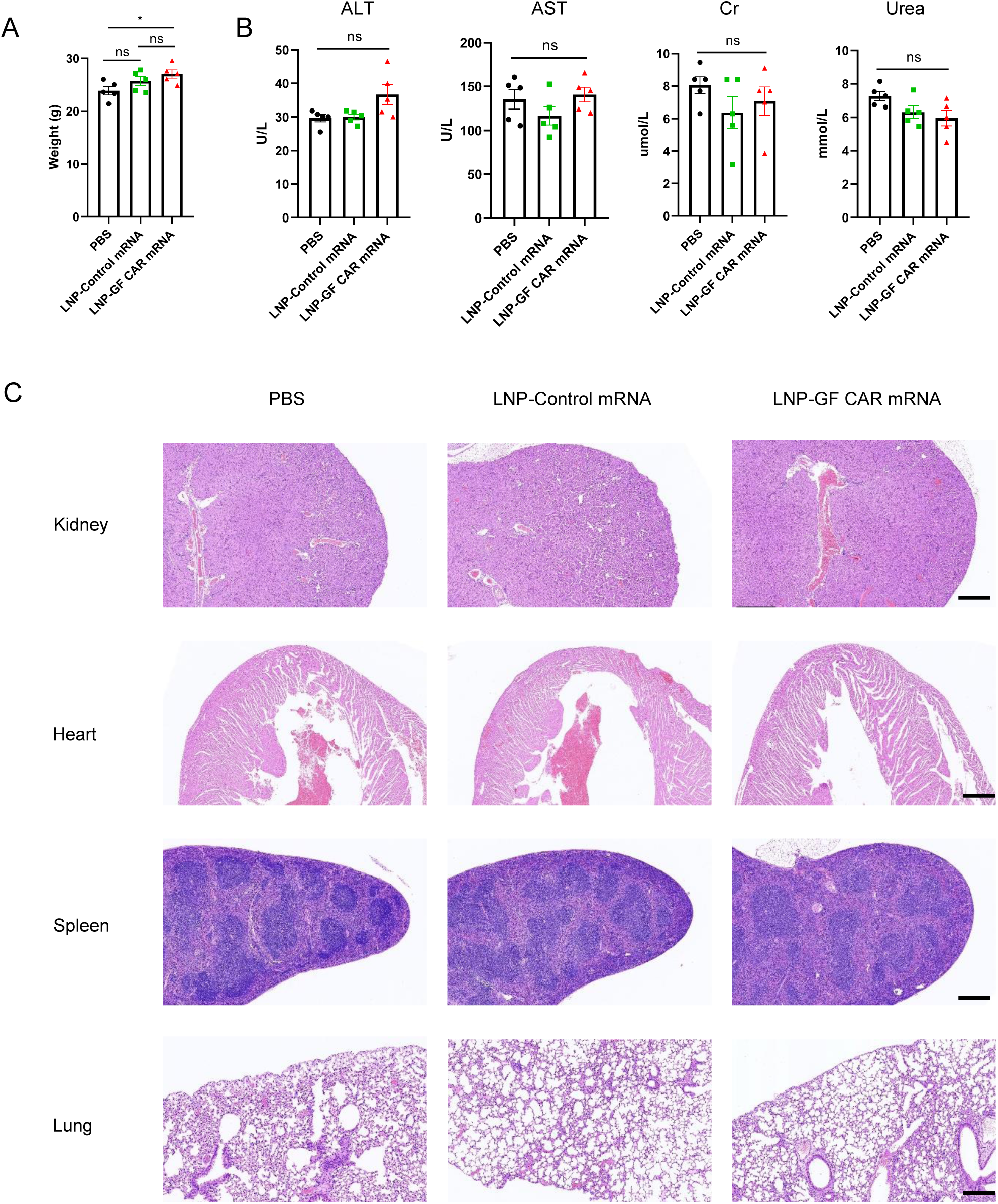
Safety evaluation of LNP-GF CAR mRNA. A: Weight of mice in each treatment group (*n* = 5). B: Serum of mice with orthotopic HCC tumors in each treatment group was collected. The levels of liver enzymes (ALT, AST), Cr, and Urea were measured (*n* = 5). C: Tissues of mice with orthotopic HCC tumors in each treatment group were collected. HE staining was performed on the kidney, heart, spleen, and lung. Scale bar (kidney, heart): 600 μm, Scale bar (spleen, lung):300um. Significance was calculated with one-way ANOVA analysis with multiple comparisons and is presented as mean ± s.e.m., ns: not significant; *p < 0.05.

**Supplementary Figure 5:**
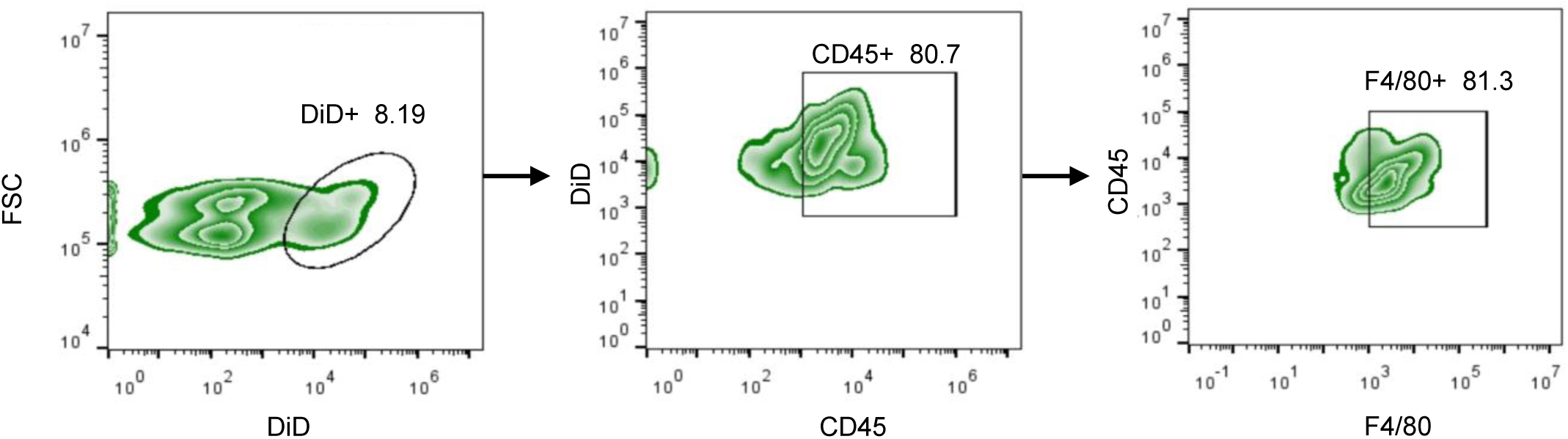
Intravenous LNP was mainly taken up by liver macrophages. A: In the preparation process of LNP, DiD dye was added at a ratio of 0.15%. DiD-labeled LNP was injected intravenically into C57 mice, and the proportion of macrophages in LNP-ingested cells in the liver was analyzed by flow cytometry 24 hours later.

**Supplementary Figure 6:**
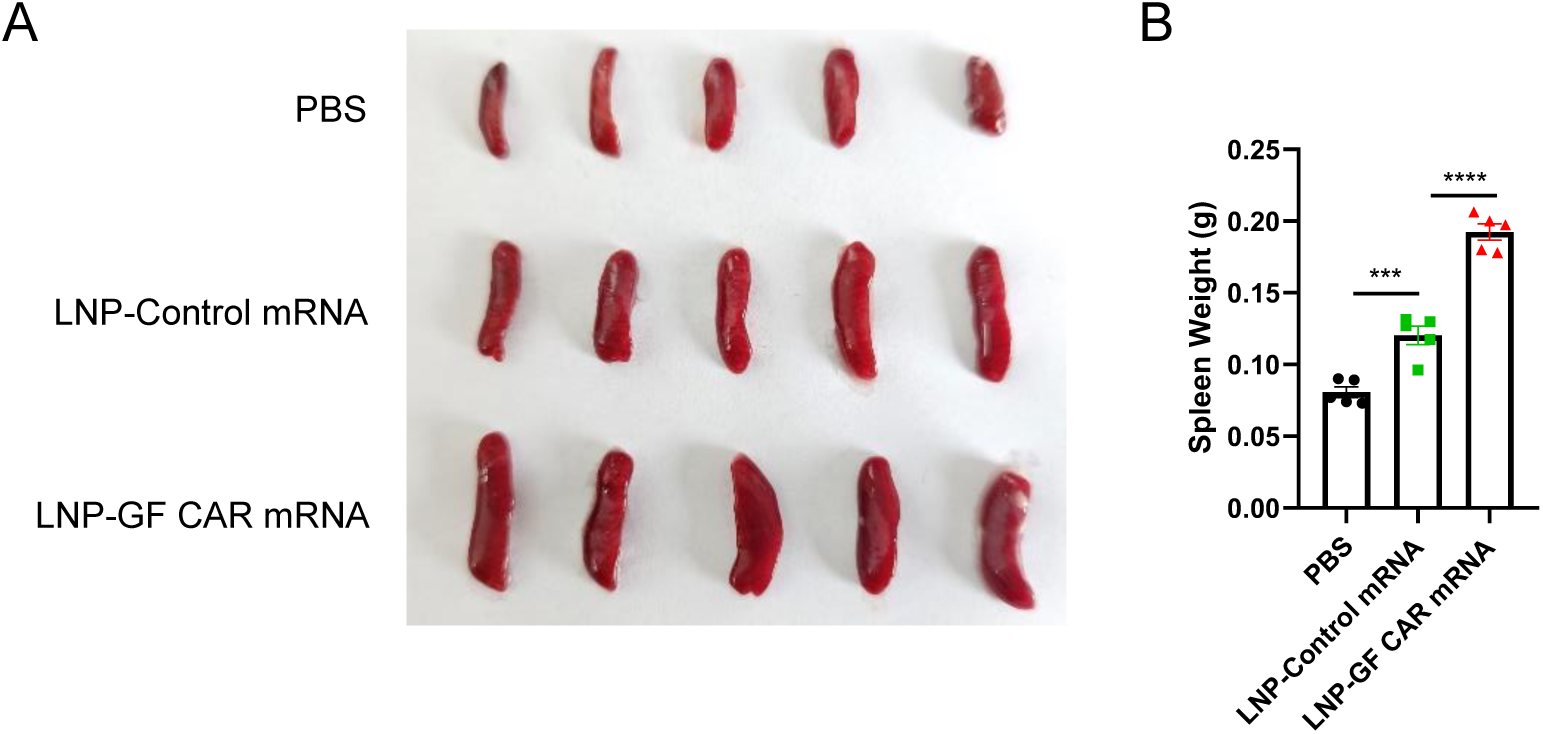
LNP-GF CAR mRNA enlarges the spleen of mice with orthotopic hepatocellular carcinoma. A, B: The spleens of mice with orthotopic HCC tumors in each treatment group were photographed (A) and weighed (B) (n=5). Significance was calculated with one-way ANOVA analysis with multiple comparisons and is presented as mean ± s.e.m., ns: not significant; ***p < 0.001, ****p < 0.0001.

**Supplementary Figure 7:**
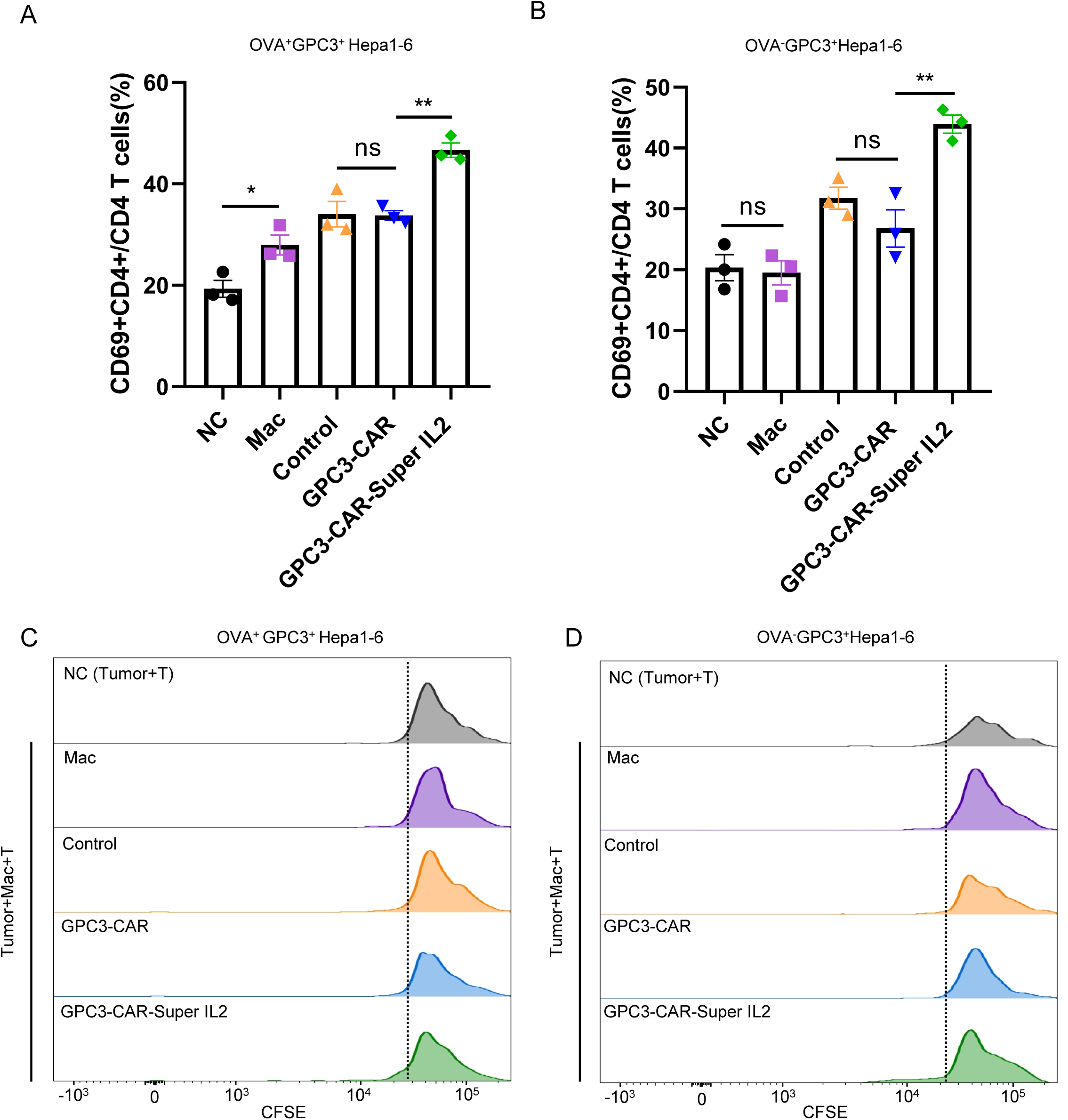
Proliferation and activation of CD4 T cells in antigen spreading assay. A-D: BMDMs were transfected with various LNPs for 24 hours and subsequently co-incubated with OVA^+^GPC3^+^Hepa1-6 or OVA^−^GPC3^+^Hepa1-6 cells and T cells derived from OT1 mice. The activation of CD4 T cells (A, B) and proliferation of CD4 T cells indicated by CFSE staining (C, D) were detected after 72 hours of co-incubation by flow cytometry. NC: No macrophages, Mac: Untransfected macrophages, Control: LNP-Control mRNA transfected macrophages, GPC3-CAR: LNP-GPC3-CAR mRNA transfected macrophages, GPC3-CAR-Super IL-2: LNP-GPC3-CAR-Super IL-2 mRNA transfected macrophages. Significance was calculated with one-way ANOVA analysis with multiple comparisons and is presented as mean ± s.e.m., ns: not significant; *p < 0.05, **p < 0.01, ***p < 0.001.

## References

1. Albelda, S.M., CAR T cell therapy for patients with solid tumors: key lessons to learn and unlearn. Nat Rev Clin Oncol, 2024. 21(1): p. 47–66.

2. Klichinsky, M., et al., Human chimeric antigen receptor macrophages for cancer immunotherapy. Nat Biotechnol, 2020. 38(8): p. 947–953.

3. Chen, K., et al., CAR-macrophage versus CAR-T for solid tumors: The race between a rising star and a superstar. Biomol Biomed, 2024. 24(3): p. 465–476.

4. Zong, Y., et al., Lipid Nanoparticle (LNP) Enables mRNA Delivery for Cancer Therapy. Adv Mater, 2023. 35(51): p. e2303261.

5. Cheng, Q., et al., Selective organ targeting (SORT) nanoparticles for tissue-specific mRNA delivery and CRISPR-Cas gene editing. Nat Nanotechnol, 2020. 15(4): p. 313–320.

6. Rurik, J.G., et al., CAR T cells produced in vivo to treat cardiac injury. Science, 2022. 375(6576): p. 91–96.

7. de Visser, K.E. and J.A. Joyce, The evolving tumor microenvironment: From cancer initiation to metastatic outgrowth. Cancer Cell, 2023. 41(3): p. 374–403.

8. Sahai, E., et al., A framework for advancing our understanding of cancer-associated fibroblasts. Nat Rev Cancer, 2020. 20(3): p. 174–186.

9. Feng, B., et al., Cancer-associated fibroblasts and resistance to anticancer therapies: status, mechanisms, and countermeasures. Cancer Cell Int, 2022. 22(1): p. 166.

10. Feig, C., et al., Targeting CXCL12 from FAP-expressing carcinoma-associated fibroblasts synergizes with anti-PD-L1 immunotherapy in pancreatic cancer. Proc Natl Acad Sci U S A, 2013. 110(50): p. 20212–7.

11. Peng, D., et al., Targeting TGF-β signal transduction for fibrosis and cancer therapy. Mol Cancer, 2022. 21(1): p. 104.

12. Batlle, E. and J. Massagué, Transforming Growth Factor-β Signaling in Immunity and Cancer. Immunity, 2019. 50(4): p. 924–940.

13. Kersten, K., et al., Spatiotemporal co-dependency between macrophages and exhausted CD8(+) T cells in cancer. Cancer Cell, 2022. 40(6): p. 624–638.e9.

14. Zhang, S., et al., SET/PP2A signaling regulates macrophage positioning in hypoxic tumor regions by amplifying chemotactic responses. Exp Mol Med, 2022. 54(10): p. 1741–1755.

15. Du, Y., et al., Potential crosstalk between SPP1 + TAMs and CD8 + exhausted T cells promotes an immunosuppressive environment in gastric metastatic cancer. J Transl Med, 2024. 22(1): p. 158.

16. Tharp, K.M., et al., Tumor-associated macrophages restrict CD8(+) T cell function through collagen deposition and metabolic reprogramming of the breast cancer microenvironment. Nat Cancer, 2024.

17. Zheng, X., et al., Glypican-3: A Novel and Promising Target for the Treatment of Hepatocellular Carcinoma. Front Oncol, 2022. 12: p. 824208.

18. Lei, A., et al., A second-generation M1-polarized CAR macrophage with antitumor efficacy. Nat Immunol, 2024. 25(1): p. 102–116.

19. Hernandez, R., et al., Engineering IL-2 for immunotherapy of autoimmunity and cancer. Nat Rev Immunol, 2022. 22(10): p. 614–628.

20. Ren, J., et al., Interleukin-2 superkines by computational design. Proc Natl Acad Sci U S A, 2022. 119(12): p. e2117401119.

21. Wu, R., et al., Comprehensive analysis of spatial architecture in primary liver cancer. Sci Adv, 2021. 7(51): p. eabg3750.

22. Wang, Y.F., et al., Spatial maps of hepatocellular carcinoma transcriptomes reveal spatial expression patterns in tumor immune microenvironment. Theranostics, 2022. 12(9): p. 4163–4180.

23. Xin, L., et al., Fibroblast Activation Protein-α as a Target in the Bench-to-Bedside Diagnosis and Treatment of Tumors: A Narrative Review. Front Oncol, 2021. 11: p. 648187.

24. Kloss, C.C., et al., Dominant-Negative TGF-β Receptor Enhances PSMA-Targeted Human CAR T Cell Proliferation And Augments Prostate Cancer Eradication. Mol Ther, 2018. 26(7): p. 1855–1866.

25. Wieser, R., et al., Signaling activity of transforming growth factor beta type II receptors lacking specific domains in the cytoplasmic region. Mol Cell Biol, 1993. 13(12): p. 7239–47.

26. Wang, T., et al., Influenza-trained mucosal-resident alveolar macrophages confer long-term antitumor immunity in the lungs. Nat Immunol, 2023. 24(3): p. 423–438.

27. Pan, K., et al., CAR race to cancer immunotherapy: from CAR T, CAR NK to CAR macrophage therapy. J Exp Clin Cancer Res, 2022. 41(1): p. 119.

28. Zhang, H., et al., Define cancer-associated fibroblasts (CAFs) in the tumor microenvironment: new opportunities in cancer immunotherapy and advances in clinical trials. Mol Cancer, 2023. 22(1): p. 159.

29. Wang, L.C., et al., Targeting fibroblast activation protein in tumor stroma with chimeric antigen receptor T cells can inhibit tumor growth and augment host immunity without severe toxicity. Cancer Immunol Res, 2014. 2(2): p. 154–66.

30. Li, F., et al., Development of Nectin4/FAP-targeted CAR-T cells secreting IL-7, CCL19, and IL-12 for malignant solid tumors. Front Immunol, 2022. 13: p. 958082.

31. Bughda, R., et al., Fibroblast Activation Protein (FAP)-Targeted CAR-T Cells: Launching an Attack on Tumor Stroma. Immunotargets Ther, 2021. 10: p. 313–323.

32. Roberts, E.W., et al., Depletion of stromal cells expressing fibroblast activation protein-α from skeletal muscle and bone marrow results in cachexia and anemia. J Exp Med, 2013. 210(6): p. 1137–51.

33. Tran, E., et al., Immune targeting of fibroblast activation protein triggers recognition of multipotent bone marrow stromal cells and cachexia. J Exp Med, 2013. 210(6): p. 1125–35.

34. Komohara, Y. and M. Takeya, CAFs and TAMs: maestros of the tumour microenvironment. J Pathol, 2017. 241(3): p. 313–315.

35. Gunaydin, G., CAFs Interacting With TAMs in Tumor Microenvironment to Enhance Tumorigenesis and Immune Evasion. Front Oncol, 2021. 11: p. 668349.

36. Tajaldini, M., et al., Cancer-associated fibroblasts (CAFs) and tumor-associated macrophages (TAMs); where do they stand in tumorigenesis and how they can change the face of cancer therapy? Eur J Pharmacol, 2022. 928: p. 175087.

37. Qi, J., et al., Single-cell and spatial analysis reveal interaction of FAP(+) fibroblasts and SPP1(+) macrophages in colorectal cancer. Nat Commun, 2022. 13(1): p. 1742.

38. Wesselhoeft, R.A., P.S. Kowalski, and D.G. Anderson, Engineering circular RNA for potent and stable translation in eukaryotic cells. Nat Commun, 2018. 9(1): p. 2629.

39. Spolski, R., P. Li, and W.J. Leonard, Biology and regulation of IL-2: from molecular mechanisms to human therapy. Nat Rev Immunol, 2018. 18(10): p. 648–659.

40. Toumi, R., et al., Autocrine and paracrine IL-2 signals collaborate to regulate distinct phases of CD8 T cell memory. Cell Rep, 2022. 39(2): p. 110632.

